# Biosynthesis and heterologous expression of cacaoidin, the first member of the lanthidin family of RiPPs

**DOI:** 10.1101/2020.05.20.105809

**Authors:** Fernando Román-Hurtado, Marina Sánchez-Hidalgo, Jesús Martín, Francisco Javier Ortíz-López, Olga Genilloud

## Abstract

Cacaoidin is the first member of the new lanthidin RiPP family, a lanthipeptide produced by the strain *Streptomyces cacaoi* CA-170360 with unprecedented features such as an unusually high number of D-amino acids, a double methylation in the N-terminal alanine and a tyrosine residue glycosylated with a disaccharide. In this work, we describe the complete identification, cloning and heterologous expression of the cacaoidin biosynthetic gene cluster, which shows unique RiPP genes.

## 2. Introduction

Actinomycetes are an extremely diverse group of Gram-positive, filamentous bacteria with high GC content genomes (1) considered as one of the most prolific sources for the discovery of new natural products (2,3). Among all the actinomycetes, the genus *Streptomyces* produces over 70-80% of the secondary metabolites with described therapeutic properties (4).

The increasing number of sequenced genomes has revealed that actinomycetes carry the genetic potential to produce many more secondary metabolites than those detected under laboratory conditions (5). The development of bioinformatic tools to identify the presence of new secondary metabolite Biosynthetic Gene Clusters (BGCs), such as antiSMASH (6) or MiBIG (7) has permitted the development of targeted genome mining strategies directed at specific families of compounds (8).

Ribosomally synthesized and Post-translationally modified Peptides (RiPPs) are a group of secondary metabolites with a large structural diversity. Most of these compounds are synthesized as a longer precursor peptide, containing an N-terminal leader peptide that usually guides secretion and is excised from the C-terminal core peptide, which finally becomes the mature RiPP (9) after undergoing a broad diversity of post-translationally modifications (PTMs).

We have recently described the discovery of the antibiotic cacaoidin, the first reported member of lanthidins, a new RIPP with unprecedented structural characteristics not found in other lanthipeptides (10) (Figure 1). This 23-amino acid molecule, produced by *Streptomyces cacaoi* CA-170360, contains several PTMs, some of them shared with other lanthipeptides. Cacaoidin presents a C-terminal amino acid S-[(Z)-2-aminovinyl-3-methyl]-D-cysteine (AviMeCys) by oxidative decarboxylation of the C-terminal cysteine. AviMeCys is formed both in lanthipeptides and linaridins (11) by LanD (12) or LinD (13) enzymesrespectively, belonging to the HFCD (Homo-oligomeric Flavin-containing Cysteine Decarboxylase) protein family (12). Cacaoidin also shows a lanthionine (Lan) ring, another characteristic of lanthipeptides (14). These thioether cross-links involve the dehydration of Ser and Thr residues to 2,3-didehydroalanine (Dha) and 2,3-didehydrobutyrine (Dhb), respectively, followed by the addition of the Cys thiol to the unsaturated amino acid. None of the linaridins described to date contains lanthionine bridges, although some show Dhb; however, no obvious homologues of lanthipeptide dehydratases are present in their BGCs (15). Cacoidin presents an unprecedented *N, N*, dimethyl lanthionine (NMe_2_Lan), typical of linaridins, a RIPP family that lacks lanthionines. The *N, N*-dimethylation is introduced by α-N-methyltransferases homologous to CypM (16, 17), but is not found in lanthipeptides BGCs. These structural features common to both families of lanthipeptides and linaridins support the proposal of cacoidin as the first reported member of the lanthidins (10).

**Figure 1.**
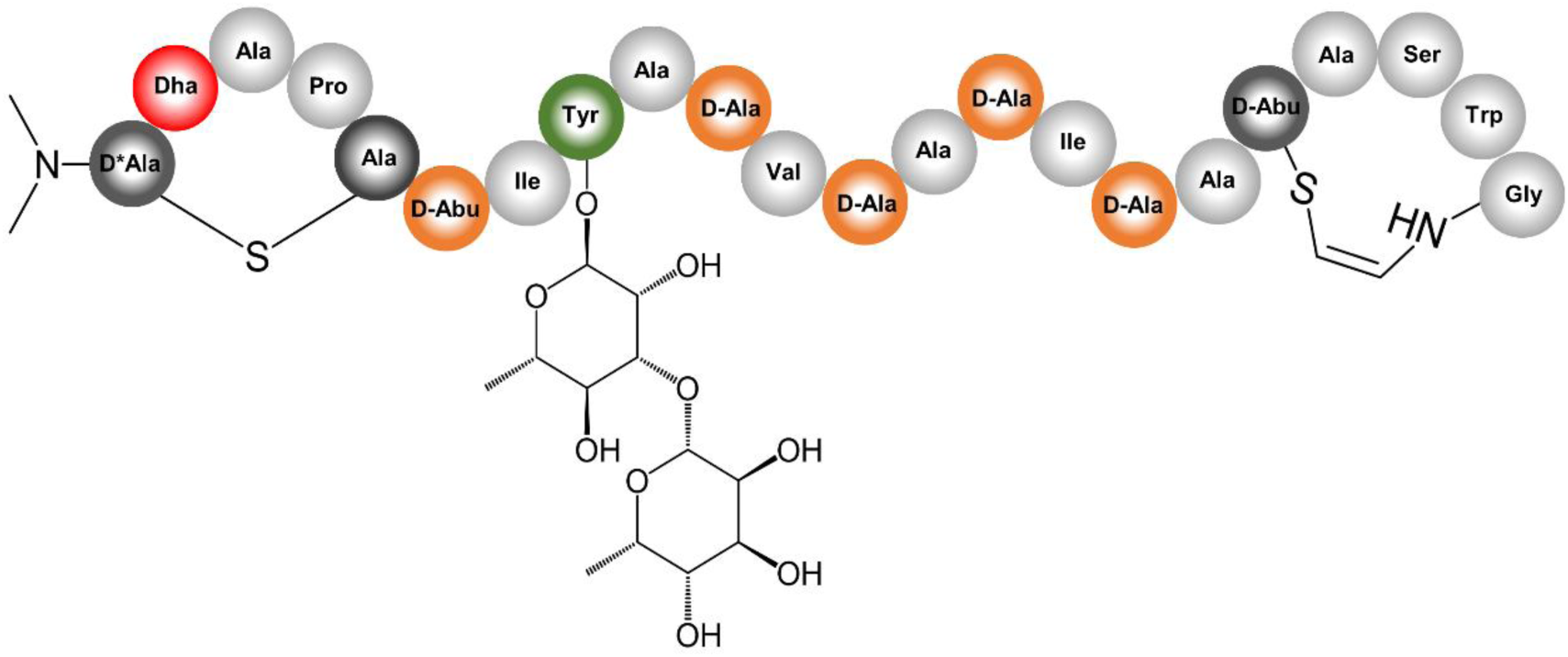
Structure of cacaoidin

Cacaoidin also presents other unusual structural features, such as a high number of D-amino acids including D-Abu and an O-glycosylated tyrosine residue carrying a non-previously reported disaccharide formed by α-L-rhamnose and β-L-6-deoxy-gulose. Cacaoidin shows potent antibacterial activity against MRSA (Methicillin resistant *Staphylococcus aureus*) (MIC 0.5 µg/mL) and moderate activity against a clinical isolate of *Clostridium difficile* (MIC 4 µg/mL) (10).

In this work, we present the identification and analysis of the cacaoidin BGC from the genome analysis of *Streptomyces cacaoi* CA-170360, showing its distinct gene cluster organization. We show that the cacaoidin BGC contains all the genes required for the antibiotic biosynthesis that was successfully produced by heterologous expression.

## 3. Results and discussion

### 3.1. Sequencing of *S. cacaoi* CA-170360 genome and identification of cacaoidin BGC

CA-170360 genome sequence was obtained with a combination of PacBio and Illumina approaches. *De novo* PacBio sequencing of CA-170360 genome provided 2 contigs of 5,971,081 bp and 2,704,105 bp, which were used as reference to map the 163 contigs obtained through Illumina sequencing, and used to correct PacBio frameshifts caused by the high GC content (73.1 %).

In order to identify the BGC responsible for the production of cacaoidin, the genome was analyzed with antiSMASH (6), BAGEL4 (18) and PRISM (19), Many BGCs were predicted, but none of these bioinformatic tools could predict the BGC responsible of cacaoidin, biosynthesis suggesting that the discovery of novel bioactive NPs by genome mining is still a challenge.

The C-terminal sequence of cacaoidin (Thr-Ala-Ser-Trp-Gly-Cys) was used as the query in a tBLASTn using the whole genome sequence to search for the gene encoding this peptide. A 162 bp Open Reading Frame (ORF) was found, which helped to elucidate the final structure of the peptide (10). Cacaoidin structural gene *caoA* encodes a 23-amino acid C-terminal core peptide (SSAPCTIYASVSASISATASWGC) following a predicted 30-amino acid N-terminal leader peptide (MGEVVEMVAGFDTYADVEELNQIAVGEAPE). Neither the leader nor the core peptide *caoA* sequences showed high sequence similarity with any other lanthipeptide or linaridin (Supporting Figure 1).

Considering the final structure of cacaoidin (10) (Figure 1) and the BLAST analysis of the ORFs located up- and downstream of *caoA*, we identified a putative 30 Kb BGC (*cao* cluster) containing 27 ORFs that were associated to the biosynthesis (Figure 2, Supporting Table 2, Supporting Table 3). Interestingly, no homologous genes of known dehydratases or cyclases commonly found in the four current classes of lanthipeptides nor in the class of linaridins could be identified in this region.

**Figure 2.**
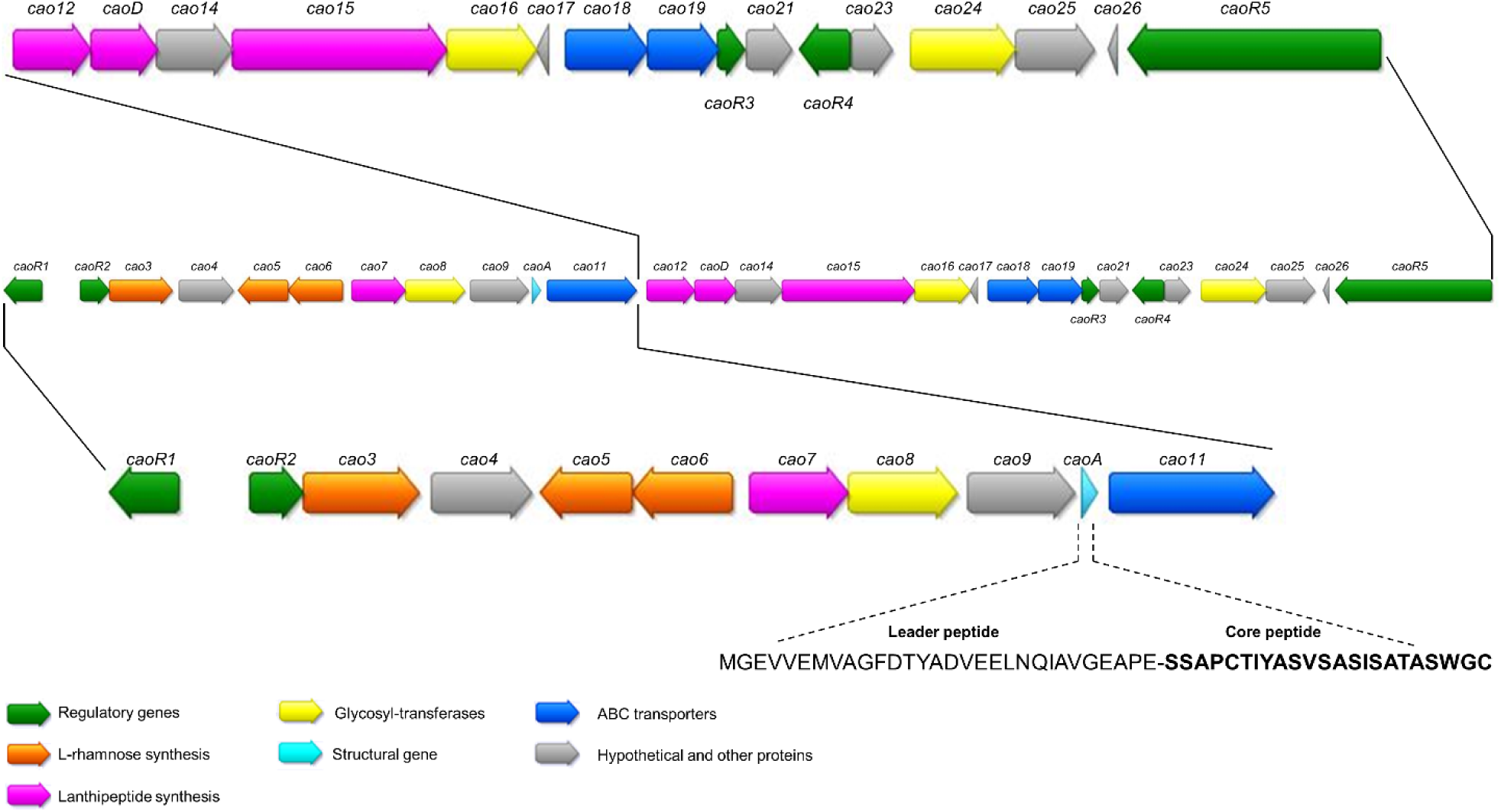
Schematic representation of the BGC of cacaoidin, where caoA codes for the precursor peptide. The sequences of the leader and core peptides of cacaoidin are shown.

The BLAST analysis (Supporting Table 2) and the secondary structure given by HHpred (Supporting Table 3) of each ORF led us to putatively assign them a role in the PTMs of cacaoidin core peptide involving the AviMeCys ring and lanthionine formation, the terminal N, N di-methylation, the incorporation of D-amino acids, the disaccharide biosynthesis and the tyrosine glycosylation.

The BGC encodes a putative cypemycin decarboxylase CypD homologue (CaoD) containing a conserved phosphopantothenoylcysteine (PPC) synthetase/decarboxylase domain. CaoD has little sequence similarity with CypD and LanD enzymes, both belonging to the HFCD protein family, and involved in the catalysis of the oxidative decarboxylation of the C-terminal cysteine residue in the presence of a flavin cofactor (20, 21). The presence of the PPC domain support a potential role in the oxidative decarboxylation and, consequently, it can be postulated that CaoD may be involved in the formation of the AviMeCys ring.

The formation of lanthionine rings is accomplished by different dehydratases and cyclases depending on the lanthipeptide class (I-IV) (15). In class I, a dehydratase (LanB) generates the Dha and Dhb and a cyclase (LanC) adds the Cys thiol. In class II, a single modification enzyme (LanM) is involved that contains an N-terminal dehydratase domain and a C-terminal LanC-like domain. In classes III and IV lanthionine rings are produced also by a single enzyme, called LanKC for class III and LanL for class IV. Both enzymes show an N-terminal phospho-Ser/phosphor-Thr lyase domain, a central kinase-like domain and a C-terminal cyclase domain which contains Zn-binding ligands only in LanL (15).

Surprisingly none of the ORFs present in the *cao* cluster showed any homology with LanC, LanM, LanKC or LanL proteins. The BLAST analysis of Cao7, which was identified as a hypothetical protein, showed some degree of homology with the N-terminal sequence of a LanC-like protein from *Raineyella antarctica* (WP_139283243.1), but Cao7 did not contain the characteristic conserved cyclase domain. Both proteins show a HopA1 conserved domain (PFAM17914), that has been described in the HopA1 effector protein from *Pseudomonas syringae* (22), that was shown to directly bind the Enhanced Disease Susceptibility 1 (EDS1) complex in *Arabidopsis thaliana*, activating the immune response signaling pathway. Future research is still needed to determine the function of this protein that can only be tentatively proposed as potential new type of lanthionine synthetase.

The N-terminal Ala dimethylation of cypemycin, the prototypical member of linaridins (11), is carried out by the S-adenosylmethionine (SAM)-dependent methyltransferase CypM (13, 16). No CypM homologues have been found in the genome of the producing strain. Within the *cao* cluster, *cao4* encodes a putative O-methyltransferase containing the conserved Methyltransf_2 domain, also belonging to the family of SAM-dependent methyltransferases. Many class I lanthipeptide clusters from actinobacteria contain an O-methyltransferase, generically known as LanS. Two types of LanS enzymes have been described: LanS_A_, which incorporates β-amino acid isoaspartate (23) and LanS_B_, which methylates the C-terminal carboxylate of a RiPP precursor (24). Cao4 shows very low homology with both types of LanS proteins. Since cacaoidin does not contain isoaspartate nor a C-terminal methylation, the role of Cao4 in the N, N-methylation is currently under study.

D-Amino acids provide a wide variety of properties to lanthipeptides, such as resistance to proteolysis, induction of bioactivity or structural conformation (25). However, only L-amino acids can be added by the ribosomal machinery, so the way to introduce D-stereocenters into lanthipeptides is modifying L-Ser and L-Thr, leading to Dha and Dhb, which will be subjected to a diastereoselective hydrogenation, to finally incorporate D-Ala and D-Abu, respectively (15, 24). This reaction is carried out by dehydrogenases generically called LanJ (15), which are divided in two classes, namely the zinc-dependent dehydrogenases (LanJ_A_) and the flavin-dependent dehydrogenases (LanJ_B_). LanJ_B_ is able to reduce both Dha and Dhb, whereas LanJ_A_ can only hydrogenate Dha. To date, only two LanJ_B_ enzymes have been characterized, CrnJ_B_ and BsjJ_B_, involved in the biosynthesis of carnolysin (26) and bicereucin (27), respectively. Recently, another flavin-dependent oxidoreductase (LahJ_B_) has been described in the putative lanthipeptide biosynthetic gene cluster *lah* (24).

Within the *cao* BGC, the protein Cao12 shows homology with LLM class flavin-dependent oxidoreductases and might be involved in the incorporation of D-amino acids.

The cacaoidin disaccharide has not previously reported and is formed by α-L-rhamnose and β-L-6-deoxy-gulose. Four proteins are required for the synthesis of α-L-rhamnose: a Glucose-1-phosphate thymidylyltransferase (RmlA), a dTDP-D-glucose 4,6-dehydratase (RmlB), a dTDP-4-keto-6-deoxy-D-glucose 3,5-epimerase (RmlC) and a dTDP-4-keto-6-deoxy-L-mannose reductase (RmlD), although the corresponding genes do not have to be necessarily clustered (Figure 3) (28). The *cao* BGC only contains three of the four genes *rmlA, rmlB* and *rmlD*. Nevertheless, a BLAST search of RmlC against CA-170360 whole genome sequence also shows the presence of a *rmlC* gene and additional *rmlA, rmlB* and *rmlD* genes outside the cacaoidin cluster.

**Figure 3.**
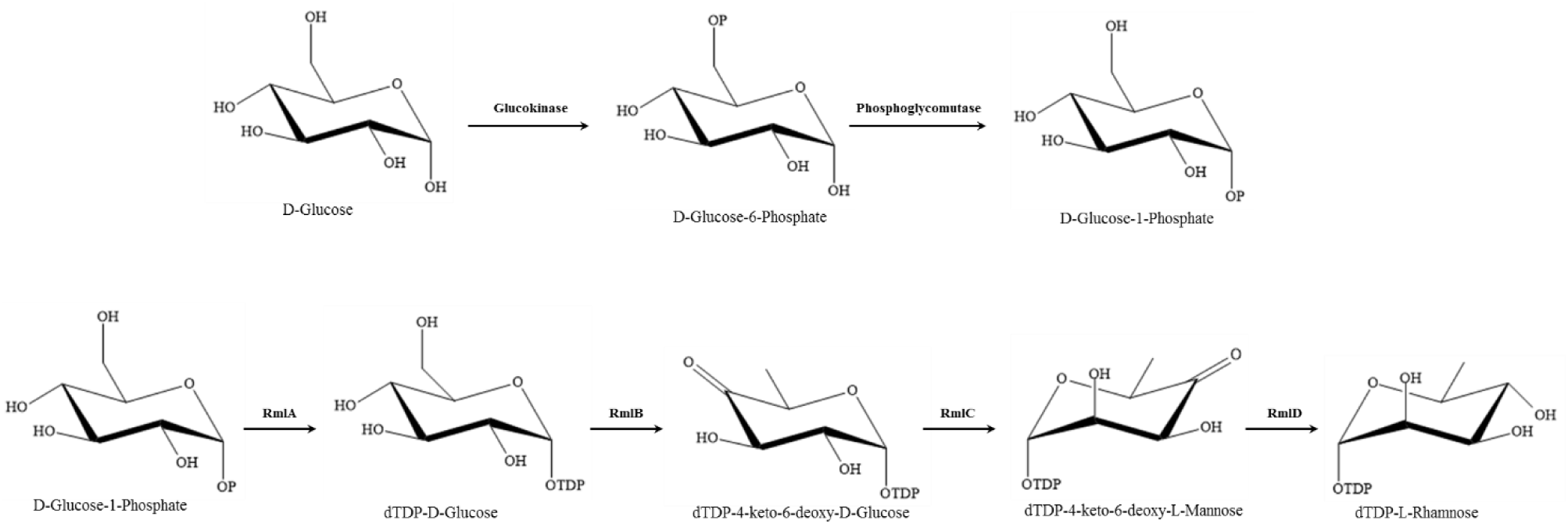
Schematic presentation of the biosynthesis of dTDP-L-Rhamnose from D-Glucose

Bleomycin, tallysomycin and zorbamycin are antitumor antibiotics which incorporate NDP-L-gulose or NDP-6-deoxy-L-gulose to their structures and their biosynthesis was used as reference to look for the presence of similar proteins being encoded in our genome. The sugar biosynthesis in the pathways of these compounds involves four classes of enzymes enzymes (Figure 4): a NTP-sugar synthase (BlmC/TlmC/ZbmC), a sugar epimerase (BlmG/TlmG), a GDP-mannose-4,6-dehydratase (ZbmL) and aNAD-dependent sugar epimerase (ZbmG) (29). Despite no homologues of these genes were found in the cacaoidin BGC, a BLAST search in the total genome sequence of CA-170360 permitted to identify some protein homologues. These include a D-glycero-beta-D-manno-heptose 1-phosphate adenylyltransferase homologous to BlmC/TlmC/ZbmC (48% similarity); a NAD-dependent epimerase/dehydratase homologous to BlmE/TlmE (38.7% similarity); a GDP-mannose 4,6-dehydratase with 62% similarity to ZbmL; and a dTDP-glucose 4,6-dehydratase with 34% similarity with ZbmG. However, as none of these homologue proteins are found associated within the same cluster, no conclusions can be made for the β-L-6-deoxy-gulose biosynthesis.

**Figure 4.**
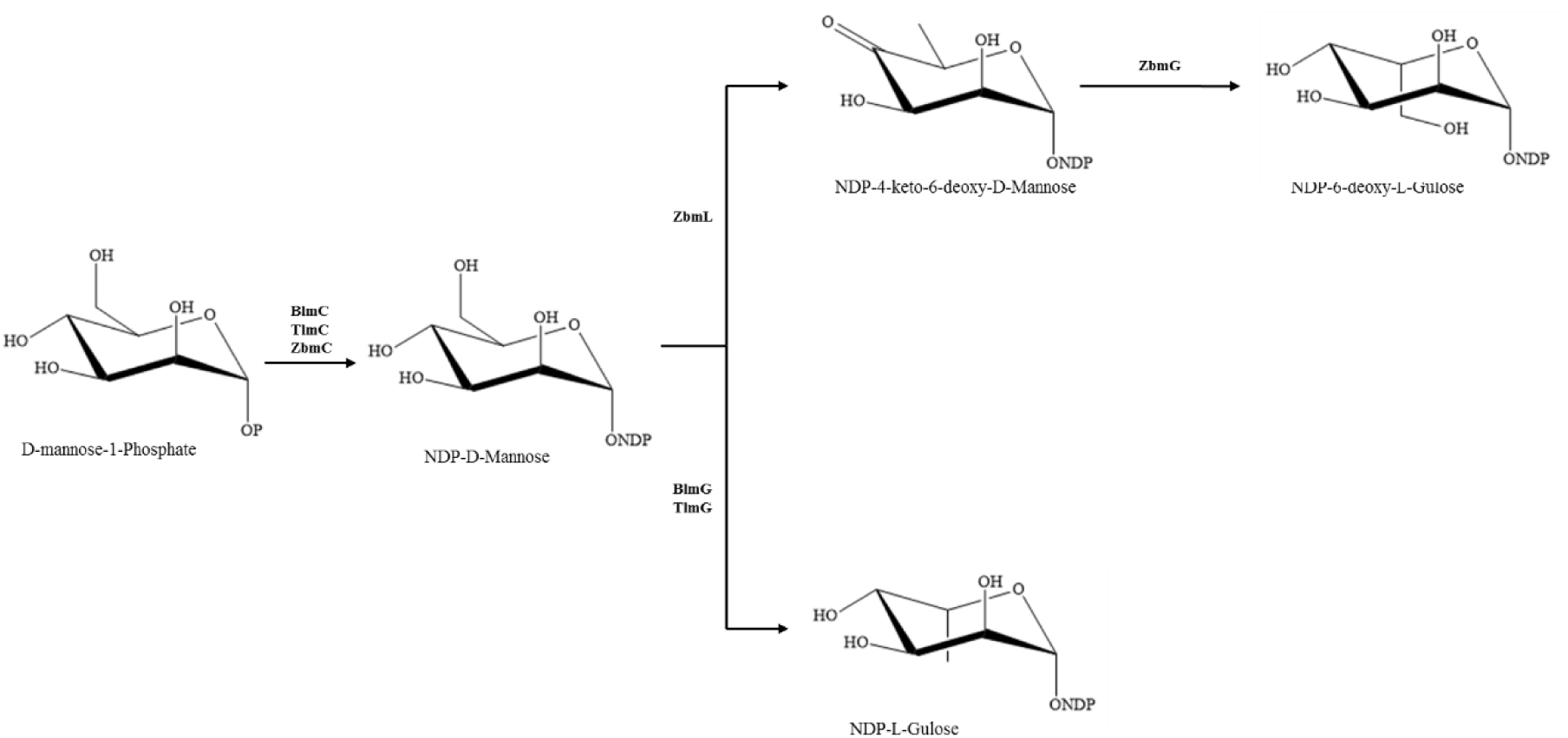
Proposed pathway for the β-L-6-deoxy-gulose sugar biosynthesis for the BLM, TLM and ZMB compounds.

As one of its unusual structural features of cacaoidin, the disaccharide α-L-rhamnose-β-L-6-deoxy-gulose is O-linked to the aromatic ring of the tyrosine residue. While asparagine N-glycosylation and serine, threonine or hydroxyproline O-glycosylation have been reported in many natural glycopeptides (30), the O-glycosylation of tyrosine is not common. Up to date, the only natural products undergoing a tyrosine O-glycosylation are the lipoglycopeptide antibiotics mannopeptimycins, produced by *Streptomyces hygroscopicus*, which contain an O-linked di-mannose (31). In prokaryotes, the O-glycosylation of tyrosine residues has been also reported in the S-layer of the cell envelope of *Paenibacillus alvei, Thermoanaerobacter thermohydrosulfuricus* and *Thermoanaerobacterium thermosaccharolyticum* strains. In *P. alvei* CCM 2051^T^, a polymeric branched polysaccharide is O-glycosidically linked *via* an adaptor to specific tyrosine residues of the S-layer protein SpaA by the O-oligosaccharyl:protein transferase WsfB (32). This protein is encoded within the *slg* cluster, that carries the genes necessary for the biosynthesis of this glycan chain. The *cao* cluster lacks an homologue of WsfB, so we cannot propose a candidate that O-glycosylates the tyrosine residue of cacaoidin.

The *cao* cluster contains three glycosyltransferases (GTs) (Cao8, Cao16, Cao24) belonging to two families of glycosyltransferases, GT-2 and GT-4. Cao8 and Cao16 belong to the family GT-2, that contain a GT_2_WfgS_like domain, involved in O-antigen biosynthesis. Cao8 and Cao16 show 42% identity (54% similarity) and 43% identity (52% similarity), respectively, with an UDP-Glc:alpha-D-GlcNAc-diphosphoundecaprenol beta-1,3-glucosyltransferase WfgD from *Streptomyces* sp. F-1, which catalyzes the addition of Glc, the second sugar moiety of the O152-antigen repeating unit, to GlcNAc-pyrophosphate-undecaprenol (33). Cao24 belong to the family GT-4 that has a GT4_GtfA-like domain and a conserved RfaB domain, involved in the cell wall and membrane biosynthesis (34).

Despite the presence of three GTs in the cacaoidin BGC, only two amino sugars are detected in the structure. The three GTs could be involved in the glycosylation as it has already been proposed for other clusters with more GT genes than amino sugars linked in the compound and proposed to work together and to be required to achieve efficient glycosylation. The biosynthesis of PM100117/PM100118 (35), saquayamycins (36) and sipanmycin (37) are some of these examples. On the basis of the absolute configurations of the cacaoidin sugar moieties, it has been proposed that Cao8 and Cao16 might work cooperatively to attach the α-L-rhamnose unit, while Cao24 would incorporate the β-L-6-deoxygulose unit (10).

Two rhamnosyltransferases (WsfF and WsfG) have been identified in the *slg* cluster of *P. alvei* as the responsible of the attatchment of the L-rhamnose to the tyrosine residue. In the case of mannopeptimycins, two peptide mannosyltransferases (MppH and MppI) would O-glycosylate the tyrosine residue. However, in all these cases low homologies were found between these enzymes and the glycosyltransferases (GTs) present in the cacaoidin cluster. Further studies are needed to confirm the role of each GT in cacaoidin biosynthesis.

Processing of leader peptide is another key step in the post-translational modification impacting in the producer immunity and transport. The N-terminal leader peptide plays a role in targeting the unmodified precursor by the posttranslational modifying enzymes, in the secretion of the peptide and in keeping the modified pre-peptide inactive (38). The enzymes responsible for the removal of the leader peptide depend on the type of lanthipeptide. Class I lanthipeptides are exported by the ABC transporter LanT and their leader peptides are cleaved by the serine protease LanP (14). In class II, both secretion and cleavage are performed by a unique enzyme with a conserved N-terminal cysteine protease domain, called LanT_P_ (39).

In the cacaoidin cluster, Cao14 encodes a putative Zn-dependent peptidase belonging to the M16 peptidase family that may be involved in the leader peptide processing. Recently, it has been reported that the leader peptide of the class III lanthipeptide NAI-112 (40) is removed by a bifunctional Zn-dependent M1-class metalloprotease, AplP, that first cleaves the N-terminal segment of the leader peptide as an endopeptidase, and subsequently removes the remaining leader sequence through its aminopeptidase activity (41). Leader peptide removal in class III lanthipeptides does not have a general mechanism. In fact, in labyrinthopeptins and curvopeptins, an endopeptidase is involved in the partial N-terminal segment removal of the leader peptide and the remaining overhang is progressively trimmed off by an additional aminopeptidase (42). In other cases, such as flavipeptin (43), a designated prolyl oligopeptidase (POP) is involved in the cleavage of the leader peptide of modified precursor peptides at the C-terminal of a Pro residue, although it is not clear if a second aminopeptidase is needed to complete the leader peptide removal. Class IV lanthipeptides often lack a designated protease to cleave the leader peptide, but it has been reported that some of them might also use AplP homologs (41). When AplP and Cao14 were compared, both proteins showed a low homology degree (17.2% identity, 25.4% similarity). Future research will clarify if Cao14 is the cacaoidin leader peptidase and if it has a dual function as endo- and aminopeptidase.

Besides, three ABC transporters were found in the pathway (Cao11, Cao18 and Cao19) that might be responsible of the export and self-resistance of cacaoidin. In addition to the active removal of the leader sequence coupled to active transport, two non-universal immunity strategies have been adopted by strains producing class I and II lanthipeptides. This active transport is mediated by the ABC type transport system LanFEG and sequestering the mature lanthipeptide in the extracellular environment by LanI immunity proteins (44). A self-immunity mechanism has not been deeply studiedfor class III and IV lanthipeptides but, as in the case of cacaoidin, it has also been proposed that ABC transporters could play a role in the self-resistance of the producer strains (45).

Gene expression in the cacaoidin cluster seems to be under the control of different classes of regulators. Five transcriptional regulators are found involving one LuxR (CaoR1), two HTH-type XRE (CaoR2 and CaoR3), one TetR (CaoR4) and one SARP (CaoR5) regulators. XRE and TetR have been described as transcriptional repressors (46, 47) while LuxR and SARP have been described as transcriptional activators (48, 49). Further studied of the regulation of lanthipeptide biosynthesis will clarify their role in the production of the antibiotic.

Among the remaining eight genes identified in the *cao* cluster, six of the proteins (Cao7, Cao14, Cao17, Cao21, Cao 25 and Cao26) do not have any defined functions. Cao9 is a phosphotransferase containing a conserved APH domain, which confers resistance to various aminoglycosides (50). It has been reported that some phosphotransferases may provide self-resistance against aminoglycosides, as shown for streptomycin 6-phosphotransferase (51) or CapP, involved in the resistance to capuramycin antibiotics (52). The role of Cao9 in the biosynthesis cluster of cacaoidin is currently unknown. A protein belonging to the START/RHO_alpha_C/PITP/Bet_v1/CoxG/CalC (SRPBCC) superfamily is also present in the cluster (Cao23). SRPBCC proteins share α/β helix-grip-fold structures and have a deep hydrophobic ligand-binding pocket (53, 54). This superfamily contains aromatase/cyclase (ARO/CYC) domains of proteins such as tetracenomycin from *Streptomyces glaucescens* (55), and the SRPBCC domains of *Streptococcus mutans* Smu.440 and related proteins (56).

The HHpred analysis of each ORF was also used for the detection of RiPP precursor peptide Recognition Elements (RREs) (57). These RRE are structurally similar conserved precursor peptide-binding domain present in the majority of known prokaryotic RiPP modifying enzymes and are usually responsible for the leader peptide recognition (57). These RREs are related to the small peptide chaperone PqqD, involved in the biosynthesis of pyrroloquinoline quinone (PQQ) (58), which reportedly binds to PqqA (precursor peptide) to do its function (59). In this analysis, we used HHPred to search PqqD-like domains in the putative biosynthetic proteins from *cao* gene cluster, even those with unknown functions. In fact, the identification of an RRE within the protease StmE, involved in lasso peptide streptomonomicin (STM) biosynthesis, and an “ocin_ThiF_like” cyclodehydratase (TOMM F) protein from TOMM (Thiazole/Oxazole-Modified Microcin) biosynthetic gene clusters, allowed to assign its non-previously proposed function. However, no RREs were found in the Cao proteins, suggesting the possibility of alternative leader peptide recognition domains that are unrelated to the already known RREs (57). As homology detection algorithms will become more accurate and more sequences will become available, additional RREs will be found.

### 3.2. Cloning and heterologous expression of cacaoidin BGC

The strain *S. cacaoi* CA-170360 is reluctant to genetic manipulation, limiting the obtention of knockdown mutants to confirm the involvement of the *cao* gene cluster in the biosynthesis of cacaoidin. To confirm that the *cao* cluster was responsible of antibiotic biosynthesis, we cloned and heterologously expressed the cacaoidin BGC in the genetically amenable host *Streptomyces albus* J1074.

We followed the CATCH method to clone a 40 Kb region containing the *cao* BGC into the pCAP01 vector (60), yielding pCAO. pCAO was introduced into NEB-10-beta *E. coli* ET12567 cells by electroporation. A triparental conjugation was carried out between *E. coli* ET12567/pCAO, *E. coli* ET12567/pUB307 and *S. albus* J1074 spores (61). Five positive transconjugants, alongside the negative control (*S. albus* J1074/pCAP01) and the wild-type strain CA-170360, were grown in R2YE for 14 days at 28°C to confirm the production of the targeted antibiotic. After acetone extraction of the cultures, organic solvent was evaporated, and the aqueous extracts in 20% DMSO were analyzed by LC-HRESI-TOF. The analysis of the extracts from pCAO transconjugants confirmed the presence of cacaoidin as peaks at 3.35 minutes were detected, coincident with the retention time of elution of cacaoidin in the wild type strain and purified cacaoidin standards. The perfect correlation between the UV spectrum, exact mass and isotopic distribution of cacaoidin standards and the components isolated from the transconjugants *S. albus* J1074/pCAO undoubtedly demonstrated that they correspond to cacaoidin (Supporting Figure 2). These preliminary results clearly confirm that the *cao* BGC cloned in pCAO is enough to ensure the biosynthesis of cacaoidin.

### 3.3. Comparison with other clusters

To study if more lanthidin-encoding clusters can be found within actinomycetes, a BLAST search against the NCBI whole genome shotgun sequences database was performed, and clusters with high degree of homology to cacaoidin were found in the strains *Streptomyces cacaoi* subsp. *cacaoi* strain NRRL B-1220 (MUBL01000486), *Streptomyces* sp. NRRL F-5053 (JOHT01000009), *Streptomyces* sp. NRRL S-1868 (JOGD01000003), *Streptomyces cacaoi* subsp. *cacaoi* NBRC 12748 (BJMM01000002.1) and *Streptomyces cacaoi* subsp. *cacaoi* OABC16 (VSKT010000024) (Figure 5, Supporting Table 4). An alignment of the precursor peptide of the cacaoidin in all homologous clusters showed that no variations in the protein sequence were found (Supporting Figure 3). No other cacaoidin-related peptides or pathways were found in the databases, indicating that the cacaoidin BGC is very conserved. A phylogenetic tree generated using neighbor-joining method and corrected with the Jukes and Cantor algorithm (62, 63) showed the close relatedness of strain *Streptomyces cacaoi* CA-170360 with the strains that also contain the *cao* cluster, which was highly supported by the bootstrap values (Supporting Figure 4). Moreover, when the 16S rDNA sequences of the strains harboring the cacaoidin BGC were analyzed in EzBiocloud, all of them were identified as *Streptomyces cacaoi* (data not shown), indicating that the cacaoidin BGC is so far limited to this specific species, with no identifiable orthologs in other species. Several genome comparative studies have found strain-specific BGCs in some species of *Streptomyces*, reflecting that chemical novelty can be found at the strain level and that the analysis of the genomes of closely related strains constitutes a promising approach for the identification of novel BGCs (64, 65).

**Figure 5.**
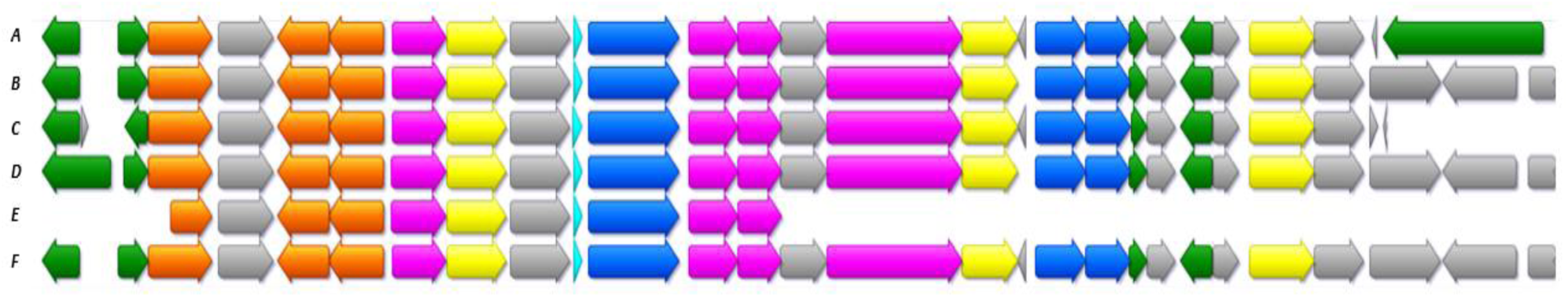
Schematic representation of the alignment of cacaoidin BGC in Streptomyces cacaoi CA-170360 and the clusters found in NCBI with high degree of homology. Most of them belong as well to a strain of Streptomyces cacaoi. A: Streptomyces cacaoi CA-170360; B: Streptomyces cacaoi NBRC 12748; C: Streptomyces sp. NRRL S-1868; D: Streptomyces sp. NRRL F-5053; E: Streptomyces cacaoi NRRL B-1220; F: Streptomyces cacaoi OABC16).

Nevertheless, the analysis of below-threshold scores of CaoA BLAST results, together with the search of HopA1 domain-containing proteins similar to Cao7, allowed us to find some pathways that could encode new lanthidins (Supporting Figure 5). The alignment of the hypothetical precursor peptides shows the presence of some conserved residues that possibly could be involved in the leader peptide recognition by biosynthetic enzymes (Supporting Figure 6). Also, the analysis of the ORFs present in all these clusters show that all of them contain a HopA1 domain-containing protein, a LLM flavin-dependent oxidoreductase, a CypD-related protein, a Zn-dependent or S9 peptidase and a putative phosphotransferase (Supplorting Figure 5). Most of these clusters also contain an O-methyltransferase. These data suggest a broader distribution of potential BGCs enconding new lanthidins. However, we will need to have more lanthidin molecules described, before we can conclude the existence of a minimal set of genes required to produce a lanthidin.

## 4. Conclusions

Cacaoidin is the first member of the new lanthidin RiPP family, characterized by structural features of lanthipeptide and linaridin families, and encoded by a new unprecedented RiPP BGC organization that could not be detected by any bioinformatic tool. The lack of homology with common lanthionine ring formation or double N-terminal dimethylation enzymes suggests an alternative mechanism of biosynthesis. The other unusual structural features of cacaoidin, such as the high number of D-amino acids or the O-glycosylation of tyrosine are supported by the presence in the *cao* cluster of protein homologues of a LLM class flavin-dependent oxidoreductase and three glycosyltransferases The heterologous expression of cacaoidin, has demonstrated that the *cao* cluster contains all the necessary genes to biosynthesize cacaoidin, and future research is needed to clarify the unassigned functions of the *cao* genes Cacaoidin BGC cluster was only found in the genomes of all *Streptomyces cacaoi* strains publicly available and not in any other species, suggesting that this cluster may be a species-specific trait. Undoubtedly, cacaoidin BGC has an unprecedented genetic organization, completely different from any other previously described RiPP cluster. Moreover, the detection of similar putative lanthidin homologous clusters opens the door to the study of a new exciting family of RiPPs.

## 5. Material and Methods

Detailed descriptions of all procedures are provided in the Supporting Information. Primer sequences for the cacaoidin gene cluster cloning are included in Supporting Table S1. The *cao* BGC sequence is available in the National Center for Biotechnology Information (NCBI) database under accession GenBank number MT210103.

## 6. Acknowledgments

This work is supported by Novo Nordisk Foundation grant NNF16OC0021746. The authors thank Daniel Oves-Costales for helpful advice during the whole process and the Microbiology and Chemistry areas of Fundación MEDINA for the technical support. We thank José Antonio Salas and the University of Oviedo for kindly provide strains *Streptomyces albus* J1074 and *Escherichia coli* ET12567/pUB307.

## References

1. Berdy, J. (2012). Thoughts and facts about antibiotics: where we are now and where we are heading. J Antibiot. 65(8), 385–395, doi: 10.1038/ja.2012.27

2. Berdy, J. (2005). Bioactive microbial metabolites. J Antibiot. 58(1), 1–26, doi: 10.1038/ja.2005.1

3. Genilloud, O. (2017). Actinomycetes: still a source of novel antibiotics. Nat. Prod. Rep. 34, 1203–1232, doi: 10.1039/C7NP00026J

4. Ventura, M., Canchaya, C., Tauch, A., Chandra, G., Fitzgerald, G. F., Chater, K. F., van Sinderen, D. (2007). Genomics of Actinobacteria: tracing the evolutionary history of an ancient phylum. Microbiol. Mol. Biol. Rev. 71, 495–548, doi: 10.1128/MMBR.00005-07

5. Gomez-Escribano, J. P., Alt, S. and Bibb, M. (2016). Next Generation Sequencing of Actinobacteria for the Discovery of Novel Natural Products. Mar. Drugs. 14(4), 78–97, doi: 10.3390/md14040078

6. Blin, K., Wolf, T., Chevrette, M. G., Lu, X., Schwalen, C. J., Kautsar, S. A., Suarez-Duran, H. G., de Los Santos, E. L. C., Kim, H. U., Nave, M., Dickschat, J. S., Mitchell, D. A., Shelest, E., Breitling, R., Takano, E., Lee, S. Y., Weber, T. and Medema, M. H. 2017) antiSMASH 4.0-improvements in chemistry prediction and gene cluster boundary identification. Nucleic Acids Res. 45(W1), W36–W41, doi: 10.1093/nar/gkx319

7. Medema, M. H., Kottmann, R., Yilmaz, P., Cummings, M., Biggins, J. B., Blin, K., de Bruijin, I., Chooi, Y. H., Claesen, J., Coates, J. et al. (2015). Minimum Information about a Biosynthetic Gene cluster. Nat. Chem. Biol. 11, 625–631. DOI: 10.1038/nchembio.1890

8. Genilloud, O. (2018). Mining Actinomycetes for Novel Antibiotics in the Omics Era: Are We Ready to Exploit This New Paradigm? Antibiotics (Basel). 7(4), 85–98, doi: 10.3390/antibiotics7040085

9. Arnison, P. G., Bibb, M. J., Bierbaum, G., Bowers, A. A., Bugni, T. S., Bulaj, G., Camarero, J. A., Campopiano, D. J., Challis, G. L., Clardy, J., et al. (2013). Ribosomally synthesized and post-translationally modified peptide natural products: overview and recommendations for a universal nomenclature. Nat. Prod. Rep. 30(1), 108–160, doi: 10.1039/C2NP20085F

10. Ortiz-López, F. J., Carretero-Molina, D., Sánchez-Hidalgo, M., Martín, J., Gónzalez, I., Román-Hurtado, F., de la Cruz, M., García-Fernández, S., Reyes, F., Deisinger, J., Schneider, T and Genilloud, O. (2020). Cacaoidin, first member of the new lanthidin RiPP family.

11. Claesen, J. and Bibb, M. (2010). Genome mining and genetic analysis of cypemycin biosynthesis reveal an unusual class of posttranslationally modified peptides. Proc. Natl. Sci. U. S. A. 107(37), 16297–16302, doi: 10.1073/pnas.1008608107

12. Sit, C. S., Yoganathan, S. and Vederas, J. C. (2011). Biosynthesis of aminovinyl-cysteine-containing peptides and its application in the production of potential drug candidates. Acc. Chem. Res. 44(4), 261–268, doi: 10.1021/ar1001395

13. Mo, T., Liu, W. Q., Ji, W., Zhao, J., Chen, T., Ding, W., Yu, S. and Zhang, Q. (2017). Biosynthetic insights into linaridin natural products from genome mining and precursor peptide mutagenesis. ACS Chem. Biol. 12(6), 1484–1488, doi: 10.1021/acschembio.7b00262

14. Zhang, Q., Yu, Y., Velásquez, J. E. and van der Donk, W. A. (2012). Evolution of lanthipeptide synthetases. Proc. Natl. Acad. Sci. U. S. A. 109, 18361–18366, doi: 10.1073/pnas.1210393109

15. Repka, L. M., Chekan, J. R., Nair, S. K. and van der Donk, W. A. (2017). Mechanistic understanding of lanthipeptide biosynthetic enzymes. Chem Rev. 117(8), 5457–5520, doi: 10.1021/acs.chemrev.6b00591

16. Claesen, J. and Bibb, M. J. (2011). Biosynthesis and regulation of grisemycin, a new member of the linaridin family of ribosomally synthesized peptides produced by *Stretpmyces griseus* IFO 13350. J. Bacteriol. 193(10), 2510–2516, doi: 10.1128/JB.00171-11

17. Rateb, M. E., Zhai, Y., Ehrner, E., Rath, C. M., Wang, X., Tabudravu, J., Ebel r., Bibb, M. Kyeremeh, K., Dorrestein, P. C., Hong, K., Jaspars, M. and Deng, H. (2015). Legonaridin, a new member of linaridin RiPP from a Ghanaian *Streptomyces* isolate. Org. Biomol. Chem. 13(37). 9585–9592, doi: 10.1039/c5ob01269d

18. van Heel, A. J., de Jong, A., Song, C., Viel, J. H., Kok, J. and Kuipers, O. P. (2018). BAGEL4: a user-friendly web server to thoroughly mine RiPPs and bacteriocins. Nucleic Acids Res. 46(W1), W278–W281, doi: 10.1093/nar/gky383

19. Skinnider, M. A., Merwin, N. J., Johnston, C.W. and Magarvey, N. A. (2017). PRISM 3: Expanded prediction of natural product chemical structures from microbial genomes. Nucleic Acids Res. 45, W49–W54, doi: 10.1093/nar/gkx320

20. Ding, W., Mo, T., Mandalapu, D. and Zhang, Q. (2018). Substrate specificity of the cypemycin decarboxylase CypD. Synth. Syst. Biotechnol. 3(3), 159–162, doi: 10.1016/j.synbio.2018.09.002

21. Sit, C. S., Yoganathan, S. and Vederas, J. C. (2011). Biosynthesis of aminovinyl-cysteine-containing peptides and its application in the production of potential drug candidates. Acc. Chem. Res. 44(4), 261–268, doi: 10.1021/ar1001395

22. Park, Y., Shin, I., & Rhee, S. (2015). Crystal structure of the effector protein HopA1 from *Pseudomonas syringae*. Journal of structural biology. 189(3), 276–280. https://doi.org/10.1016/j.jsb.2015.02.002

23. Acedo, J. Z., Bothwell, I. R., An, L., Trouth, A., Frazier, C. and van der Donk, W. (2019). O-Methyltransferase-Mediated Incorporation of a β-Amino Acid in Lanthipeptides. J. Am. Chem. Soc. 141(42), 16790–16801, doi: 10.1021/jacs.9b07396

24. Huo, L., Zhao, X., Acedo, J. Z., Estrada, P., Nair, S. K. and van der Donk, W. A. (2019). Characterization of a dehydratase and methyltransferase in the biosynthesis of a ribosomally-synthesized and post-translationally modified peptide in Lachnospiraceae. ChemBioChem. 21(1-2), 190–199, doi: 10.1002/cbic.201900483

25. Yang, X. and van der Donk, W. A. (2015). Post-translational Introduction of D-Alanine into Ribosomally Synthesized Peptides by the Dehydroalanine Reductase NpnJ. J. Am. Chem. Soc. 137(39), 12426–12429, doi: 10.1021/jacs.5b05207

26. Lohans, C. T., Li, J. L. and Vederas, J. C. (2014). Structure and Biosynthesis of Carnolysin, a Homologue of Enterococcal Cytolysin with D-Amino Acids. J. Am. Chem. Soc. 136(38), 13150–13153, doi: 10.1021/ja5070813

27. Huo, L. and van der Donk, W. A. (2016). Discovery and Characterization of Bicereucin, an Unusual D-Amino Acid-Containing Mixed Two-component Lantibiotic. J. Am. Chem. Soc. 138(16), 5254–5257, doi: 10.1021/jacs.6b02513

28. Girauld, M. F. and Naismith, J. H. (2000). The rhamnose pathway. Curr. Opin. Struct. Biol. 19(6), 687–696, doi: 10.1016/s0959-440x(00)00145-7

29. Galm, U., Wendt-Pienkowski, E., Wang, L., Huang, S. X., Unsin, C., Tao, M., Coughlin, J. M. and Shen, B. (2011). Comparative analysis of the biosythetic gene clusters and pathways for three structurally related antitumor antibiotics: bleomycin, tallosomycin and zorbamycin. J. Nat. Prod. 74(3), 526–536, doi: 10.1021/np1008152

30. Varki, A., Cummings, R., Esko, J., Freeze, H., Hart, G. and Marth, J. (1999). Essentials of Glycobiology. Cold Spring Harbor (NY): Cold Spring Harbor Laboratory Press.

31. He, H., Williamson, R. T., Shen, B., Graziani, E. I., Yang, H. Y., Sakya, S. M., Petersen, P. J. and Carter, G. T. (2002). Mannopeptimycins, novel antibacterial glycopeptides from Streptomyces hygroscopicus, LL-AC98. J. Am. Chem. Soc. 124(33), 9729–9736, doi: 10.1021/ja020257s

32. Messner, P., Christian, R., Neuninger, C. and Schulz, G. (1995). Similarity of “core” structures in two different glycans of tyrosine-linked eubacterial S-layer glycoproteins. J Bacteriol. 177, 2188–2193, doi: 10.1128/jb.177.8.2188-2193.1995

33. Brockhausen, I., Hu, B., Liu, B., Lau, K., Szarek, W. A., Wang, L. and Feng, L. (2008). Characterization of Two β-1,3-Glucosyltransferases from *Escherichia coli* Serotypes O56 and O152. J. Bacteriol. 190(14), 4922–4932, doi: 10.1128/JB.00160-08

34. Pradel, E., Parker, C. T. and Schnaitman, C. A. (1992). Structures of the rfaB, rfaI, rfaJ, and rfaS genes of Escherichia coli K-12 and their roles in assembly of the lipopolysaccharide core. J Bacteriol. 174(14), 4736–4745, doi: 10.1128/jb.174.14.4736-4745.1992

35. Salcedo, R. G., Olano, C., Fernández, R., Braña, A. F., Méndez, C., de la Calle, F. and Salas, J. A. (2016). Elucidation of the glycosylation steps during biosynthesis of antitumor macrolides PM100117 and PM100118 and engineering for novel derivatives. Microb. Cell Fact. 15,187, doi: 10.1186/s12934-016-0591-7.

36. Salem, S. M., Weidenbach, S. and Rohr, J. (2017). Two cooperative glycosyltransferases are responsible for the sugar diversity of saquayamycins isolated from Streptomyces sp. KY 40-1. ACS Chem. Biol. 12, 2529–2534, doi: 10.1021/acschembio.7b00453.

37. Malmierca, M. G., Pérez-Victoria, I., Martín, J., Reyes, F., Méndez, C., Olano, C. and Salas, J. A. (2018). Cooperative involvement of glycosyltransferases in the transfer of amino sugars during the biosymthesis of the macrolactam sipanmycin by Streptomyces sp. Strain CS149. Appl. Environ. Microbiol. 84(18), doi: 10.1128/AEM.01462-18

38. Plat, A., Kluskens, L. D., Kuipers, A., Rink, R. and Moll, G. N. (2011). Requirements of the engineered leader peptide of nisin for inducing modification, export, and cleavage. Appl. Environ. Microbiol. 77, 604–611, doi: 10.1128/AEM.01503-10

39. Knerr, P. J. and van der Donk, W. A. (2012). Discovery, biosynthesis, and engineering of lantipeptides. Annu. Rev. Biochem.81, 479–505, doi: 10.1146/annurev-biochem-060110-113521

40. Addlagatta, A., Gay, L. and Matthews, B. W. (2006). Structure of aminopeptidase N from Escherichia coli suggests a compartmentalized, gated active site. Proc. Natl. Acad. Sci. U. S. A. 103(36), 13339–13344, doi: 10.1073/pnas.0606167103

41. Chen, S., Xu, B., Chen, E., Wang, J., Lu, J., Donadio, S., Ge, H. and Wang H. (2019). Zn-dependent bifunctional proteases are responsible for leader peptide processing of class III lanthipeptides. Proc. Natl. Acad. Sci. U. S. A. 116(7), 2533–2538, doi: 10.1073/pnas.1815594116

42. Krawczyk, J. M., Völler, G. H., Krawczyk, B., Kretz, J., Brönstrup, M. and Süssmuth, R. D. (2013). Heterologous expression and engineering studies of labyrinthopeptins, class III lantibiotics from Actinomadura namibiensis. Chem. Biol. 20(1), 111–122, doi: 10.1016/j.chembiol.2012.10.023

43. Völler, G. H., Krawczyk, B., Ensle, P. and Süssmuth, R.D. (2013). Involvement and unusual substrate specificity of a prolyl oligopeptidase in class III lanthipeptide maturation. J. Am. Chem. Soc. 135(20), 7426–7429, doi: 10.1021/ja402296m

44. Geiger, C., Korn, S. M., Häsler, M., Peetz, O., Martin, J., Kötter, P., Morgner, N. and Entian, K. D. (2019). LanI-Mediated Lantibiotic Immunity in Bacillus subtilis: Functional Analysis. Appl. Environ. Microbiol. 85(11), e00534–19, doi: 10.1128/AEM.00534-19

45. Méndez, C. and Salas, J. A. (2001). The role of ABC transporters in antibiotics-producing organisms: drug secretion and resistance mechanisms. Res. Microbiol. 152(2001), 341–350, doi: 10.1016/s0923-2508(01)01205-0

46. Wood, H. E., Devine, K. M. and McConnell, D. J. (1990). Characterisation of a repressor gene (xre) and a temperature-sensitive allele from the Bacillus subtilis prophage, PBSX. Gene. 96(1), 83–88, doi: 10.1016/0378-1119(90)90344-q

47. Cuthbertson, L. and Nodwell, J. R. (2013). The TetR family of regulators. Microbiol. Mol. Biol. Rev. 77(3), 440–475, doi: 10.1128/MMBR.00018-13

48. Chen, J. and Xie, J. (2011). Role and regulation of bacterial LuxR-like regulators. J. Cell Biochem. 112(10), 2694–2702, doi: 10.1002/jcb.23219

49. Li, Y., Kong, L., Shen, J., Wang, Q., Liu, Q., Yang, W., Deng, Z. and You, D. (2019). Charactetization of the positive SARP family regulator PieR for improving piericidin A1 production in Streptomyces piomogeues var. Hangzhouwanensis. Synthetic and Systems Biotechnol. 4(1), 16–24, doi: 10.1016/j.synbio.2018.12.002

50. Wright, G. D. and Thompson, P. R. (1999). Aminoglycoside phosphotransferases: proteins, structure and mechanism. Front. Biosci. 4, D9–21, doi: 10.2741/wright

51. Sugiyama, M. and Nimi, O. (1990). Streptomycin biosynthesis and self-resistance mechanism in streptomycin-producing Streptomyces griseus. Actinomycetologica. 4(1), 15–22, doi: 10.3209/saj.4_15

52. Yang, Z., Funabashi, M., Nonaka, K., Hosobuchi, M., Shibata, T., Pahari, P. and Van Lanen S. G. (2010). Functional and kinetic analysis of the phosphotransferase CapP conferring selective self-resistance to capuramycin antibiotics. J. Biol. Chem. 285(17), 12899–12905, doi: 10.1074/jbc.M110.104141

53. Radauer, C., Lackner, P. and Breiteneder, H. (2008). The Bet v1 fold: an ancient, versatile scaffold for binding of large, hydrophobic ligands. BMC Evol. Biol. 8, 286, doi: 10.1186/1471-2148-8-286

54. Iyer, L. M., Koonin, E. V. and Aravind, L. (2001). Adaptations of the helix-grip fold for ligand binding and catalysis in the START domain superfamily. Proteins. 43, 134–144, doi: 10.1002/1097-0134(20010501)43:2<134::AID-PROT1025>3.0.CO;2-I

55. Ames, B. d., Korman, T. P., Zhang, W., Smith, P., Vu, T., Tang, Y and Tsai, S. C. (2008). Crystal structure and functional analysis of tetracenomycin ARO/CYC: Implications for cyclization specificity of aromatic polyketides. National Academy of Sciences. 105(14), 5349–5354; DOI: 10.1073/pnas.0709223105

56. Nan, J., Brostromer, E., Liu, X. Y., Kristensen, O. and Su, X. D. (2009). Bioinformatics and Structural Characterization of a Hypothetical Protein from Streptococcus mutans: Implication of Antibiotic Resistance. DOI: 10.1371/journal.pone.0007245

57. Burkhart, B. J., Hudson, G. A., Dunbar, K. L. and Mitchell, D. A. (2015). A Prevalent Peptide-Binding Domain Guides Ribosomal Natural Product Biosynthesis. Nat Chem Biol. 11(8), 564–570. DOI: 0.1038/nchembio.1856

58. Klinman, J. P. and Bonnot, F. (2014). Intrigues and intricacies of the biosynthetic pathways for the enzymatic quinocofactors: PQQ, TTQ, CTQ, TPQ, and LTQ. Chem Rev. 114, 4343–4365. DOI: 10.1021/cr400475g

59. Latham, J. A., Iavarone, A. T., Barr, I., Juthani, P. V. and Klinman, J. P. (2015). PqqD is a novel peptide chaperone that forms a ternary complex with the radical S-adenosylmethionine protein PqqE in the pyrroloquinoline quinone biosynthetic pathway. J Biol Chem. 290, 12908–12918. DOI: 10.1074/jbc.M115.646521

60. Jiang, W., Zhao, X., Gabrieli, T., Lou, C., Ebenstein, Y. and Zhu, T. F. (2015). Cas9-assisted targeting of chromosome segments CATCH enables one-step targeted cloning of large gene clusters. Nat. Commun. 6, 8101. DOI: 10.1038/ncomms9101

61. Chater, K. F. and Wilde, L. C. (1980). Streptomyces albus G mutants defective in the SalGI restriction-modification system. J. Gen. Microbiol. 116, 323–334, doi:10.1099/00221287-116-2-323

62. Jukes, T. H. and Cantor, C. Evolution of protein molecules. In Mammalian Protein Metabolism; Munro, H. N., Allison, J. B., Eds.; Academic Press: New York, NY, USA, 1969; pp. 121–132.

63. Saitou, N. and Nei, M. (1987). The neighbor-joining method: A new method for reconstructing phylogenetic trees. Mol. Biol. Evol. 4, 406–425, doi: 10.1093/oxfordjournals.molbev.a040454

64. Vicente, C. M., Thibessard, A., Lorenzi, J. N., Benhadj, M., Hôtel, L., Gacemi-Kirane, D., Lespinet, O., Leblond, P and Aigle, B. (2018). Comparative genomics among closely related *Streptomyces* strains revealed specialized metabolite biosynthetic gene cluster diversity. Antibiotics (Basel). 7, 86, doi: 10.3390/antibiotics7040086

65. Seipke, R. F. (2015). Strain-level diversity of secondary metabolism in Streptomyces albus. PLoS ONE. 10, e0116457, doi: 10.1371/journal.pone.0116457

