## Supporting Material for "Biosynthesis and heterologous expression of cacaoidin, the first member of the lanthidin family of RiPPs"

**SUPPORTING ON-LINE INFORMATION**

### Material and Methods

#### Strains and plasmids

The strain *Streptomyces cacaoi* CA-170360, from Fundación MEDINA's Culture Collection, was isolated from the rhizosphere of *Brownanthus corallinus*, in the region of Namaqualand (South Africa). Electrocompetent NEB 10-β *E. coli* (New England BioLabs, Ipswich, MA, USA), *E. coli* ET12567 (LGC Standards, Manchester, NH, USA) and *E. coli* ET12567/pUB307 (kindly provided by Jose Antonio Salas) were used for plasmid transformation and intergeneric triparental conjugation. *Streptomyces albus* J1074 (1), also kindly provided by Jose Antonio Salas, was employed as heterologous expression host.

Vector pCAP01, a yeast-*E. coli*-actinobacteria shuttle vector that can integrate cloned gene clusters into the genome of heterologous actinobacteria hosts thanks to its site-specific φC31 integrase (2), was used for the cloning of the cacaoidin BGC and was a gift from Bradley Moore (Addgene plasmid #59981; http://n2t.net/addgene:59981; RRID: Addgene_59981).

#### DNA extraction and genome sequencing

Genomic DNA of the strain *Streptomyces cacaoi* CA-170360 was extracted and purified as Kieser et al described (3) from cultures on ATCC-2 liquid medium (soluble starch 20 g/L, glucose 10 g/L, NZ Amine Type E 5 g/L, meat extract 3 g/L, peptone 5 g/L, yeast extract 5 g/L, calcium carbonate 1 g/L, pH 7) grown on an orbital shaker at 28 ºC, 220 rpm and 70% relative humidity.

The genome of CA-170360 was fully sequenced *de novo*, assembled and annotated by Macrogen (Seoul, Korea; http://www.macrogen.com/), using a combined strategy of Illumina HiSeq 2500 and PacBio RSII platforms.

**Identification of cacaoidin BGC**

The sequence of the CA-170360 genome was analyzed with antiSMASH v4.2.0 (4), BAGEL4 (5), PRISM (6) and RIPPMiner (7) in order to find cacaoidin BGC. BLAST (Basic Local Alignment Search Tool) (8) and HHpred based on profile hidden Markov model (HMM) comparisons (9) weres also employed to predict the function of each gene in the biosynthesis of the lanthipeptide.

**Cas9-Assisted Targeting of Chromosome (CATCH) cloning of cacaoidin BGC**

Cloning of cacaoidin BGC was performed by CATCH. The CRISPR-Cas9 endonuclease is guided by an RNA template to cleave specific DNA targets enabling to isolate larger BGC than other techniques such as transformation-associated recombination or single-strand overlapping annealing (10).

The CATCH cloning was performed as Jian and Zhu described (11). First, we designed the 20 nt targeted sequences flanking the BGC with the CRISPy-web tool (http://crispy.secondarymetabolites.org/). This computational tool design guide RNAs close to a PAM (Protospacer-Adjacent Motif) sequence 'NGG' (12) and we performed an overlapping PCR with the Q5 High-Fidelity polymerase from New England BioLabs (Ipswich, MA, USA), and used one specific primer within the 20 nt targeted sequence (X-sgRNA-P) and two universal primers (sgRNA-F and sgRNA-R). Then, we obtained the sgRNA using the HiScribe T7 Quick Yield RNA synthesis kit (New England Biolabs). We also performed a PCR using the pCAP01 vector as backbone with primers containing a 20 nt sequence that anneals to the template plasmid and a 30 nt overhang overlapping with both ends of the BGC and this PCR product was treated with DpnI (New England BioLabs) to cleave the template plasmid.  Primers used for CATCH cloning are described in the Supporting Table 1.

*S. cacaoi* CA-170360 was cultured in ATCC-2 for 2 days on an orbital shaker at 28 ºC, 220 rpm and 70% relative humidity and the bacterial cells were embedded in low-melting agarose plugs to make the subsequent in-gel digestion with Cas9 more feasible. The plugs were treated with lysozyme, proteinase K and washing buffers, and the in-gel Cas9 digestion was performed by mixing two agarose plugs, a cleavage buffer (100 mM HEPES pH 7.5, 750 mM KCl, 0.5 mM EDTA pH 8, 50 mM MgCl2, DEPC-treated water), the sgRNAs and the Cas9 nuclease from *S. pyogenes*

(New England BioLabs) and incubating at 37 °C for 2 h. After this time the agarose plugs were melted and digested with gelase and the cleavaged DNA was precipitated with ethanol and resuspended in DNase-free water.

Then, the Cas9-digested DNA was cloned into the vector pCAP01 by Gibson Assembly using a 2x Gibson Assembly Master Mix (New England BioLabs) and incubating at 50 °C for 1 h. After the ligation, this Gibson product (pCAO) was transformed into electrocompetent NEB-10-beta *E. coli* cells. These cells were incubated in 1 mL LB broth Miller (Sigma) (37 ºC, 250 rpm) without antibiotics, and plated on Difco LB agar Lennox (37 ºC, overnight, static) containing kanamycin (50 µg/mL).

The colonies obtained on these LB agar plates were validated by digestion with HindIII/NdeI and XbaI/EcoRV restriction endonucleases from New England BioLabs.

**Heterologous expression of cacaoidin BGC**

Construction pCAO was introduced into *Streptomyces albus* J1074 host by triparental conjugation (1). First, *E. coli* ET12567 cells were electroporated with 1 µL of the construction and plated on selective LB agar plates with 50 µg/mL kanamycin. *E. coli* ET12567/pUB307 cells were also streaked on selective LB agar plates with 50 µg/mL kanamycin and 25 µg/mL chloramphenicol, which were incubated at 37 °C overnight. Then, *E. coli* ET12567/pCAO and ET12567/pUB307 cells were collected at an O.D. of 0.4-0.6, washed three times with LB liquid medium without antibiotics supplementation, suspended with 100 µL LB liquid medium and mixed with 50 µL of previously activated spores (incubated at 50 °C for 10 min) of *S. albus* J1074. These mixtures were plated onto MA plates and incubated at 28 °C overnight (around 16 h). On the next day, the conjugation MA plates were overlayed with 1.5 mL of sterilized Milli-Q water containing 50 µg/mL kanamycin and 25 µg/mL nalidixic acid. These plates were incubated again at 28 °C for 3-5 days to let the exconjugants grow. After this time of incubation, five exconjugants from each *Streptomyces* heterologous host were picked and spreaded out on MA plates containing nalidixic acid (25 µg/mL) and kanamycin (50 µg/mL).

Five plugs from these MA plates of the recombinant strain *S. albus* J1074/pCAO were used to seed 10 mL ATCC-2 tubes which were incubated at 28 °C for 2-3 days, 220 rpm, 70% humidity, and these cultures, together with the corresponding negative controls harboring empty pCAP01 vector, were used to inoculate 10 mL of R2YE (103 g/L sucrose, 0.25 g /L K2SO4, 10.12 g/L MgCl2-6H2O, 10 g/L glucose, 0.1 g/L Difco Casaminoacids, 0.05 g/L KH2PO4, 2.944 g/L CaCl2-2H2O, 3 g/L L-proline, 5.73 g/L TES buffer, 5 g/L Difco yeast extract, 5 mL 1N NaOH, 2 mL trace elements), KM4 (4 g/L glucose, 4 g/L yeast extract, 10 g/L malt extract, 2 g/L CaCO3) (12) and FR23 (5 g/L glucose, 30 g/L soluble starch from potato, 20 g/L cottonseed flour, 20 g/L cane molasses, pH 7) fermentation media. These fermentations were incubated at 28 °C, 220 rpm, 70% humidity for 20 days.

#### Extraction and detection of cacaoidin

Cultures of the recombinant strain *S. albus* J1074/pCAO, together with the negative control harboring empty pCAP01 vector, were subjected to extraction with acetone 1:1 under continuous shaking at 220 rpm for 2 hours. Then the organic solvent with the secondary metabolites extracted was separated from the biomass by centrifugation, the acetone was removed by a stream of nitrogen overnight and the extracts were resuspended to a final ratio of 20% DMSO/water. The resulting microbial extracts were filtered and analyzed by LC-HRESI-TOF.

### Supporting Tables

| Oligonucleotide | Sequence (5'-3') |
|---|---|
| Glyco1-sgRNA | TAATACGACTCACTATAGGACGACTCACGTGTCAAAGAGTTTTAGAGCTAGAAATAGCAA |
| Glyco2-sgRNA | TAATACGACTCACTATAGGGGCGAGATGCCATTCCAAGGTTTTAGAGCTAGAAATAGCAA |
| sgRNA-F | GTTTTAGAGCTAGAAATAGCAAGTTAAAATAAGGCTAGTC |
| sgRNA-R | AAAAGCACCGACTCGGTGCCACTTTTTCAAGTTGATAACGGACTAGCCTTATTTTAACT |
| pCAP01-Glyco-F | CGTGCGGTGGACCGCGCCGTGACCCCCTTGTCGAGACTTGAGGTACCTGT |
| pCAP01-Glyco-R | GGGCCGGGTTCCAGCCGGTGATGCCGTCTTTCGAGGTTACTAGTCGATCT |

Supporting Table 1. Primers used for the cloning of the *cao* biosynthetic gene cluster into the pCAP01 vector through the CATCH method

| | Closest BLAST homolog | % Identity | % Similarity | Conserved domains | Possible function |
|---|---|---|---|---|---|
| *caoR1* | DNA-binding response regulator [Streptomyces cacaoi subsp. cacaoi] | 100 | 100 | CitB (NarL/FixJ family, contains REC and HTH domains); HTH_LUXR | Positive regulator |
| *caoR2* | Helix-turn-helix domain-containing protein [Streptomyces cacaoi] | 98.44 | 100 | HTH_XRE superfamily | Negative regulator |
| *cao3* | Hypothetical protein SCA03_05120 [Streptomyces cacaoi subsp. cacaoi] | 97.81 | 100 | RmlD_sub_bind | L-rhamnose synthesis |
| *cao4* | Methyltransferase [Streptomyces cacaoi] | 100 | 100 | Methyltrans_2 (O-methyltransferase domain) | Dimethylation of N-terminal Ala |
| *cao5* | GDP-mannose 4,6-dehydratase [Streptomyces cacaoi] | 98.16 | 100 | dTDP_GD_SDR-e (dDTP-D-glucose 4,6-dehydratase) | L-rhamnose synthesis |
| *cao6* | MULTISPECIES: glucose-1-phosphate thymidylyltransferase [Streptomyces] | 100 | 100 | rmlA_long (glucose-1-phosphate thymidylyltransferase) | L-rhamnose synthesis |
| *cao7* | Hypothetical protein [Streptomyces cacaoi] | 100 | 100 | HopA1 superfamily | Unknown |
| *cao8* | Glycosyltransferase family 2 protein [Streptomyces sp. NRRL S-1868] | 99.48 | 100 | Glycos_transf_2 | Glycosylation |
| *cao9* | Phosphotransferase [Streptomyces sp. NRRL S-1868] | 100 | 100 | PKc_like (protein kinase catalytic domain) | Unknown |
| *caoA* | Hypothetical protein SCA03_05190 [Streptomyces cacaoi subsp. cacaoi] | 100 | 100 | No putative conserved domains detected | Structural gene |
| *cao11* | ABC transporter ATP-binding protein [Streptomyces sp. NRRL F-5053] | 99.66 | 100 | MdlB (ABC-type multidrug transport system, ATPase and permease component) | Lanthipeptide biosynthesis |
| *cao12* | MULTISPECIES: LLM class flavin-dependent oxidoreductase [Streptomyces] | 100 | 100 | SsuD (Flavin-dependent oxidoreductase) | Lanthipeptide biosynthesis |
| *caoD* | Hypothetical protein SCA03_05220 [Streptomyces cacaoi subsp. cacaoi] | 99.63 | 100 | PRK05579 (bifunctional phosphopantothenoylcysteine decarboxylase/phosphopantothenate synthase) | AviMeCys biosynthesis |
| *cao14* | MULTISPECIES: hypothetical protein [unclassified Streptomyces] | 99.34 | 100 | No putative conserved domains detected | Unknown |
| *cao15* | Hypothetical protein [Streptomyces sp. NHF165] | 99,42 | 100 | PqqL (predicted Zn-dependent peptidase) | Leader peptide cleavage |
| *cao16* | Glycosyltransferase family 2 protein [Streptomyces cacaoi] | 100 | 100 | Glycos_transf_2 | Glycosylation |
| *cao17* | Hypothetical protein [Streptomyces cacaoi] | 100 | 91.7 | No putative conserved domains detected | Unknown |
| *cao18* | ABC transporter ATP-binding protein [Streptomyces sp. NRRL F-5053] | 100 | 100 | CcmA (ABC-type multidrug transport system) | Lanthipeptide biosynthesis |
| *cao19* | MULTISPECIES: ABC transporter permease [Streptomyces] | 99.65 | 100 | ABC2_membrane_3 (ABC-2 family transporter protein) | Lanthipeptide biosynthesis |
| *caoR3* | Hypothetical protein SCA03_05280 [Streptomyces cacaoi subsp. cacaoi] | 100 | 100 | HTH_XRE domain | Negative regulator |

| | | | | | |
|---|---|---|---|---|---|
| *cao21* | Hypothetical protein [Streptomyces sp. NRRL S-1868] | 100 | 100 | No putative conserved domains detected | Unknown |
| *caoR4* | TetR/AcrR family transcriptional regulator [Streptomyces cacaoi] | 100 | 100 | AcrR (DNA-binding transcriptional regulator) | Negative regulator |
| *cao23* | Hypothetical protein SCA03_05310 [Streptomyces cacaoi subsp. cacaoi] | 99.39 | 100 | SRPBCC superfamily | Unknown |
| *cao24* | MULTISPECIES: glycosyltransferase family 4 protein [Streptomyces] | 100 | 100 | GT4_AmsD-like | Glycosylation |
| *cao25* | Hypothetical protein [Streptomyces cacaoi] | 99.06 | 100 | No putative conserved domains detected | Unknown |
| *cao26* | No homologues found | - | - | - | Unknown |
| *caoR5* | Tetratricopeptide repeat protein [Streptomyces sp. NRRL S-1868] | 99.31 | 100 | BTAD (Bacterial Transcriptional Activation) | Positive regulator |

*Supporting Table 2. Closest BLAST homolog for each ORF in cacaoidin BGC.*

| | Closest HHpred homolog | Probability | E-value |
|---|---|---|---|
| *caoR1* | Response regulator protein VraR; Enterococcus faecium, LiaR, response regulator | 99.96 | 1.5e-25 |
| *caoR2* | Methylphosphonate synthase | 98.06 | 0.00017 |
| *cao3* | DTDP-4-dehydrorhamnose reductase, rfbD ortholog | 99.97 | 8e-28 |
| *cao4* | O-methyltransferase family protein | 100 | 6.2e-37 |
| *cao5* | Adenosylhomocysteinase (E.C.3.3.1.1, 1.8.1.7 | 100 | 1.9e-37 |
| *cao6* | Bifunctional protein Glm | 100 | 7.8e-37 |
| *cao7* | Type III effector HopA1 | 100 | 3.4e-39 |
| *cao8* | Putative glycosyltransferase protein | 99.94 | 2.4e-25 |
| *cao9* | 5-methylthioribose kinase (E.C.2.7.1.100) | 99.9 | 2.3e-21 |
| *caoA* | Myocyte-specific enhancer factor 2A | 25.67 | 67 |
| *cao11* | Lipid A export ATP-binding/permease protein | 100 | 1.3e-89 |
| *cao12* | Bacterial luciferase, 1,2-ethanediol; monooxygenase, flavoprotein | 100 | 1e-41 |
| *caoD* | MrsD protein | 100 | 7.5e-32 |
| *cao14* | Spectinomycin phosphotransferase; protein kinase, aminoglycoside phosphotransferase, antibiotic; | 99.76 | 5.4e-17 |
| *cao15* | Insulinase family protein; Protease, M16 Family, Processing Protease | 100 | 3e-65 |
| *Por lcao16* | Glycosyltransferase | 99.95 | 3.5e-26 |
| *cao17* | Light-harvesting protein B-875 alpha chain | 54.44 | 10 |
| *cao18* | ABC transporter ATP-binding protein | 100 | 2.5e-51 |
| *cao19* | ATP-binding cassette sub-family G member | 99.93 | 7.2e-23 |
| *caoR3* | Putative transposon-related DNA-binding protein | 99.56 | 8.8e-14 |
| *cao21* | Ubiquinol-cytochrome c reductase iron-sulfur subunit | 59.39 | 56 |
| *caoR4* | TRANSCRIPTIONAL REGULATORY PROTEIN (PROBABLY TETR-FAMILY); tetR family of transcriptional regulator | 99.94 | 3.3e-24 |
| *cao23* | Conserved protein; Structural genomics, unknown function, ligand; | 99.89 | 7.2e-20 |
| *cao24* | Glycosyl transferase (E.C.2.4.1.57) | 100 | 5.8e-38 |
| *cao25* | TarM; Glycosyltransferase, GT-A, Wall teichoic acid | 97.73 | 0.0025 |
| *cao26* | ABC transporter ATP-binding protein (E.C.3.6.3.-) | 87.92 | 1.5 |
| *caoR5* | tetratricopeptide repeat sensor PH0952 | 100 | 1.2e-36 |

*Supporting Table 3. Closest HHpred  homolog for each ORF in cacaoidin BGC. HHPred probabilities give the most relevant representation of significance with ≥90% usually being considered a true positive.*

| | | *Streptomyces cacaoi* NBRC 12748 | | *Streptomyces* sp. NRRL F-5053 | | *Streptomyces* sp. NRRL S-1868 | | *Streptomyces cacaoi* NRRL B-1220 | | *Streptomyces cacaoi* OABC16 | |
|---|---|---|---|---|---|---|---|---|---|---|---|
| | | % Identity | % Similarity | % Identity | % Similarity | % Identity | % Similarity | % Identity | % Similarity | % Identity | % Similarity |
| *caoR1* | LuxR regulator | 100 | 100 | 99.2 | 99.6 | 100 | 100 | - | - | 99.6 | 100 |
| *caoR2* | XRE regulator | 97.9 | 99 | 98.1 | 98.1 | 34.4 | 40.6 | - | - | 97.9 | 99 |
| *cao3* | dTDP-4-dehydrorhamnose reductase | 97.8 | 98.1 | 99 | 99 | 96.6 | 97.1 | 99.6 | 100 | 98.1 | 98.3 |
| *cao4* | O-methyltransferase | 99.1 | 99.7 | 99.7 | 100 | 99.1 | 99.7 | 99.1 | 99.7 | 100 | 100 |
| *cao5* | dTDP-glucose 4,6-dehydratase | 97.9 | 98.8 | 97.5 | 98.2 | 98.2 | 98.8 | 97.9 | 98.8 | 98.2 | 98.8 |
| *cao6* | Glucose-1-phosphate thymidylyltransferase | 100 | 100 | 99.7 | 100 | 100 | 100 | 100 | 100 | 99.2 | 99.7 |
| *cao7* | LanM | 99.7 | 99.7 | 98.9 | 99.4 | 99.1 | 99.4 | 99.4 | 99.7 | 99.7 | 100 |
| *cao8* | Glycosyl transferase family 2 | 98.7 | 99.5 | 99.5 | 99.5 | 99.5 | 99.7 | 98.7 | 99.5 | 99.2 | 99.5 |
| *cao9* | Phosphotransferase | 99.7 | 99.7 | 99.7 | 100 | 100 | 100 | 99.7 | 99.7 | 99.5 | 100 |
| *caoA* | Cacaoidin precursor peptide | 100 | 100 | 100 | 100 | 100 | 100 | 100 | 100 | 100 | 100 |
| *cao11* | ABC transporter | 99.1 | 100 | 99.7 | 100 | 99.1 | 100 | 99.1 | 100 | 99.5 | 100 |
| *cao12* | LanJB | 99.7 | 100 | 100 | 100 | 99.7 | 100 | 99.7 | 100 | 99.7 | 100 |
| *caoD* | LanD | 99.6 | 100 | 92.1 | 92.1 | 93.3 | 93.3 | 99.6 | 100 | 94.6 | 95.7 |
| *cao14* | Hypothetical protein | 99.3 | 99.3 | 99.3 | 99.3 | 98.7 | 99 | - | - | 99 | 99 |
| *cao15* | Zn-dependent peptidase | 98.6 | 99 | 98.6 | 98.8 | 99 | 99.3 | - | - | 98.4 | 98.8 |
| *cao16* | Glycosyl transferase family 2 | 100 | 100 | 99.7 | 99.7 | 99.4 | 99.4 | - | - | 100 | 100 |
| *cao17* | Hypothetical protein | - | - | - | - | 100 | 100 | - | - | 100 | 100 |
| *cao18* | ABC transporter | 99.1 | 99.1 | 100 | 100 | 99.7 | 99.7 | - | - | 99.4 | 99.4 |
| *cao19* | ABC transporter | 99.7 | 99.7 | 99.7 | 99.7 | 99.7 | 99.7 | - | - | 99.3 | 99.7 |
| *caoR3* | XRE regulator | 100 | 100 | 100 | 100 | 99 | 100 | - | - | 99.1 | 100 |
| *cao21* | Hypothetical protein | 99.5 | 99.5 | 99.5 | 99.5 | 100 | 100 | - | - | 98.9 | 99.5 |
| *caoR4* | TetR regulator | 100 | 100 | 100 | 100 | 100 | 100 | - | - | 100 | 100 |
| *cao23* | SRPBCC | 99.4 | 99.4 | 100 | 100 | 98.8 | 98.8 | - | - | 99.4 | 100 |
| *cao24* | Glycosyl transferase family 4 | 100 | 100 | 100 | 100 | 100 | 100 | - | - | 100 | 100 |
| *cao25* | Hypothetical protein | 99.1 | 99.1 | 96.9 | 97.8 | 98.8 | 99.1 | - | - | 98.4 | 98.8 |

| | | | | | | | | | | | |
|---|---|---|---|---|---|---|---|---|---|---|---|
| *cao26* | Hypothetical protein | - | - | - | - | - | - | - | - | - | - |
| *caoR5* | SARP regulator | - | - | - | - | - | - | | - | | - |

*Supporting Table 4. BLAST homology between the ORFs in cacaoidin BGC and NCBI found clusters.*

### Supporting Figures

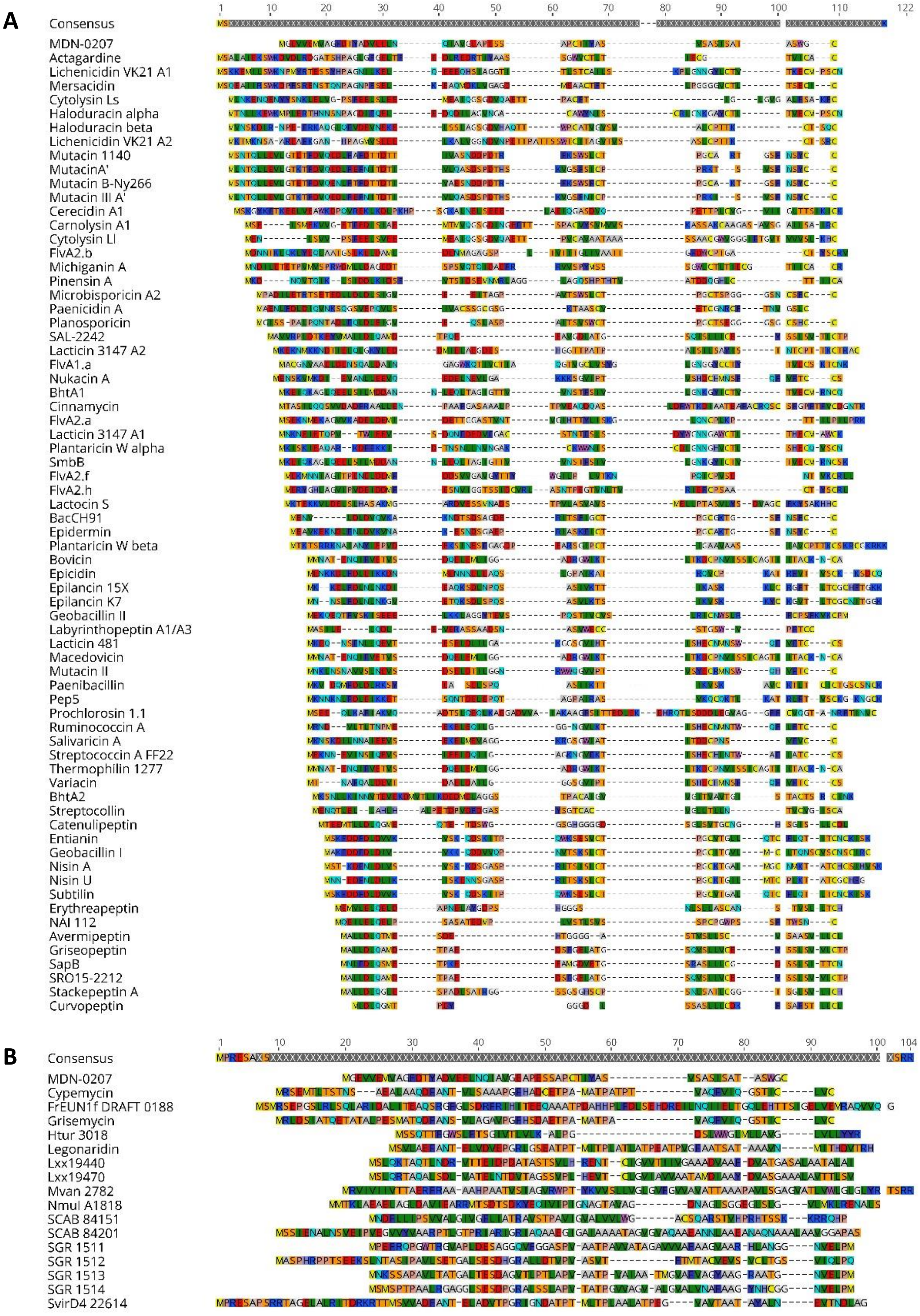

Supporting Figure 1. Alignment the structural amino acidic sequence (leader peptide + core) of cacaoidin (MDN-0207) with the sequences of other already known lanthipeptides (A) and linaridins (B).

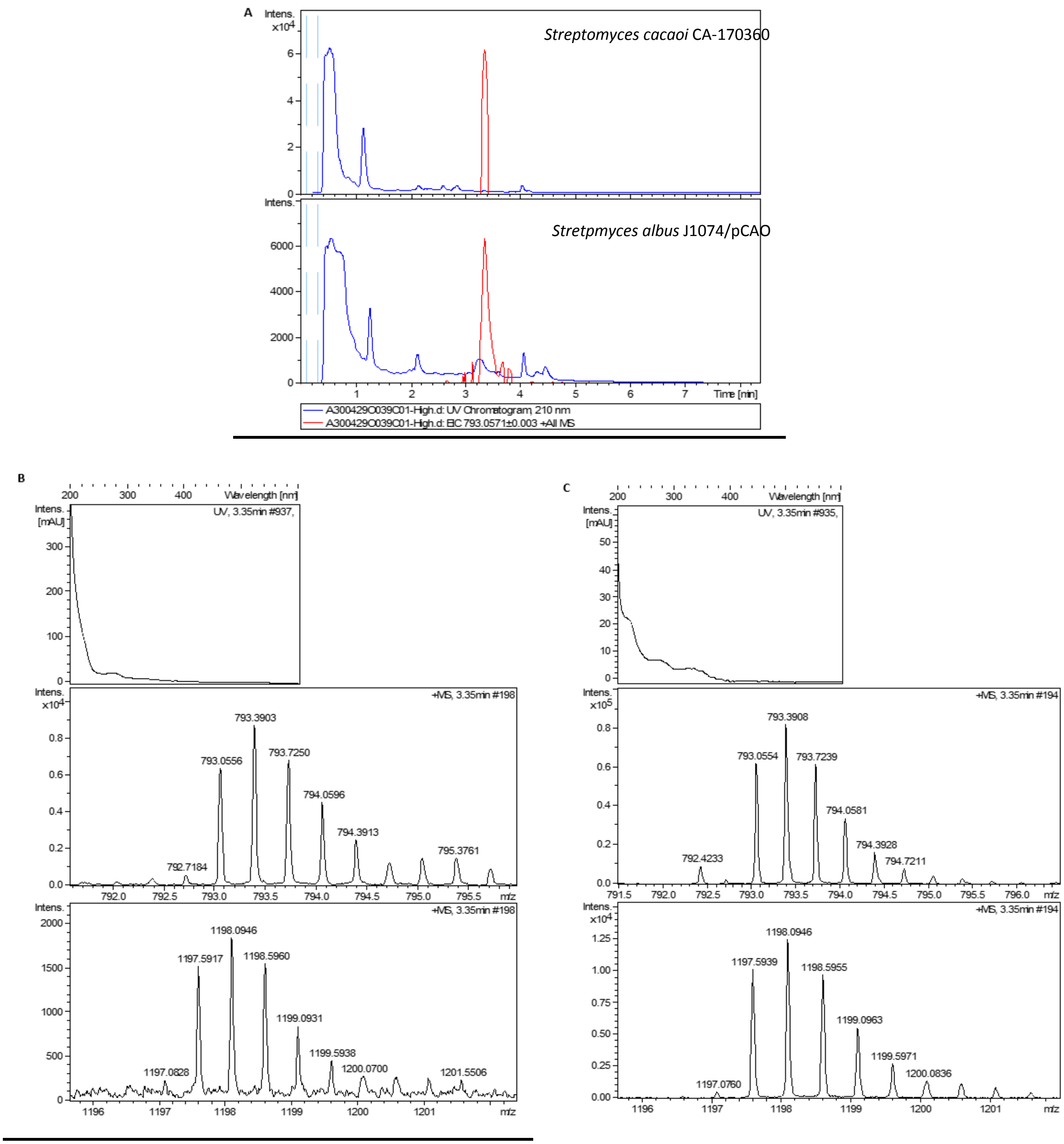


*Supporting Figure 2. (A) Chromatograms of UV absorbance at 210 nm (blue trace) and extracted ion m/z = 793.0571 ± 0.003, $C_{107}H_{162}N_{24}O_{32}S_2+NH_4^++2H^+$ of cacaoidin (MDN-0207) from original producing strain Streptomyces cacaoi CA-170360 and the heterologous producing strain Streptomyces albus J1074 with pCAO (pCAP01 + cao cluster). (B) Experimental UV and positive mass spectra from $C_{107}H_{162}N_{24}O_{32}S_2+NH_4^++2H^+$ and $C_{107}H_{162}N_{24}O_{32}S_2+2NH_4^+$ adducts from heterologous producing strain. (C) Experimental UV and positive mass spectra from $C_{107}H_{162}N_{24}O_{32}S_2+NH_4^++2H^+$ (top) and $C_{107}H_{162}N_{24}O_{32}S_2+2NH_4^+$ (bottom) adducts from original producing strain CA-170360.*

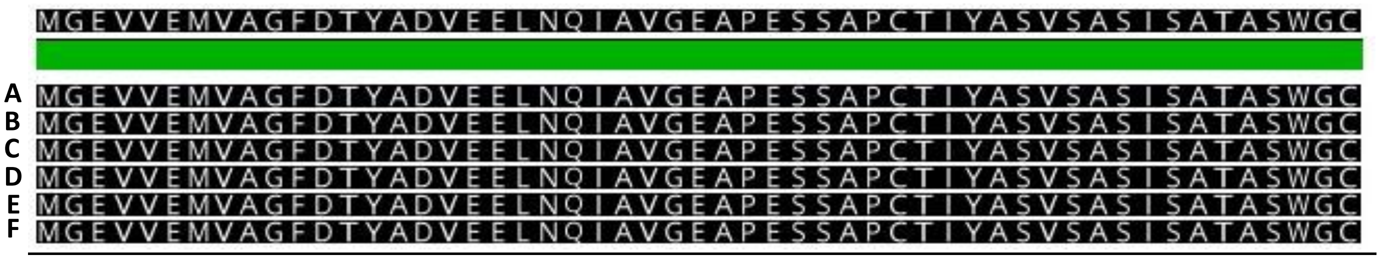


*Supporting Figure 3. Alignment of the precursor peptide of the cacaoidin in all NCBI found homologous clusters. Sequence A: precursor peptide of the* Streptomyces cacaoi *CA-170360 cluster; Sequence B: precursor peptide of the* Streptomyces cacaoi *NBRC 12748 cluster; Sequence C :precursor peptide of the* Streptomyces *sp. NRRL F-5053 cluster; Sequence D: precursor peptide of the* Streptomyces *sp. NRRL S-1868 cluster; Sequence E: precursor peptide of the* Streptomyces cacaoi *NRRL B-1220; Sequence F: precursor peptide of the* Streptomyces cacaoi *OABC16 cluster. No variations in the protein sequence are detected.*

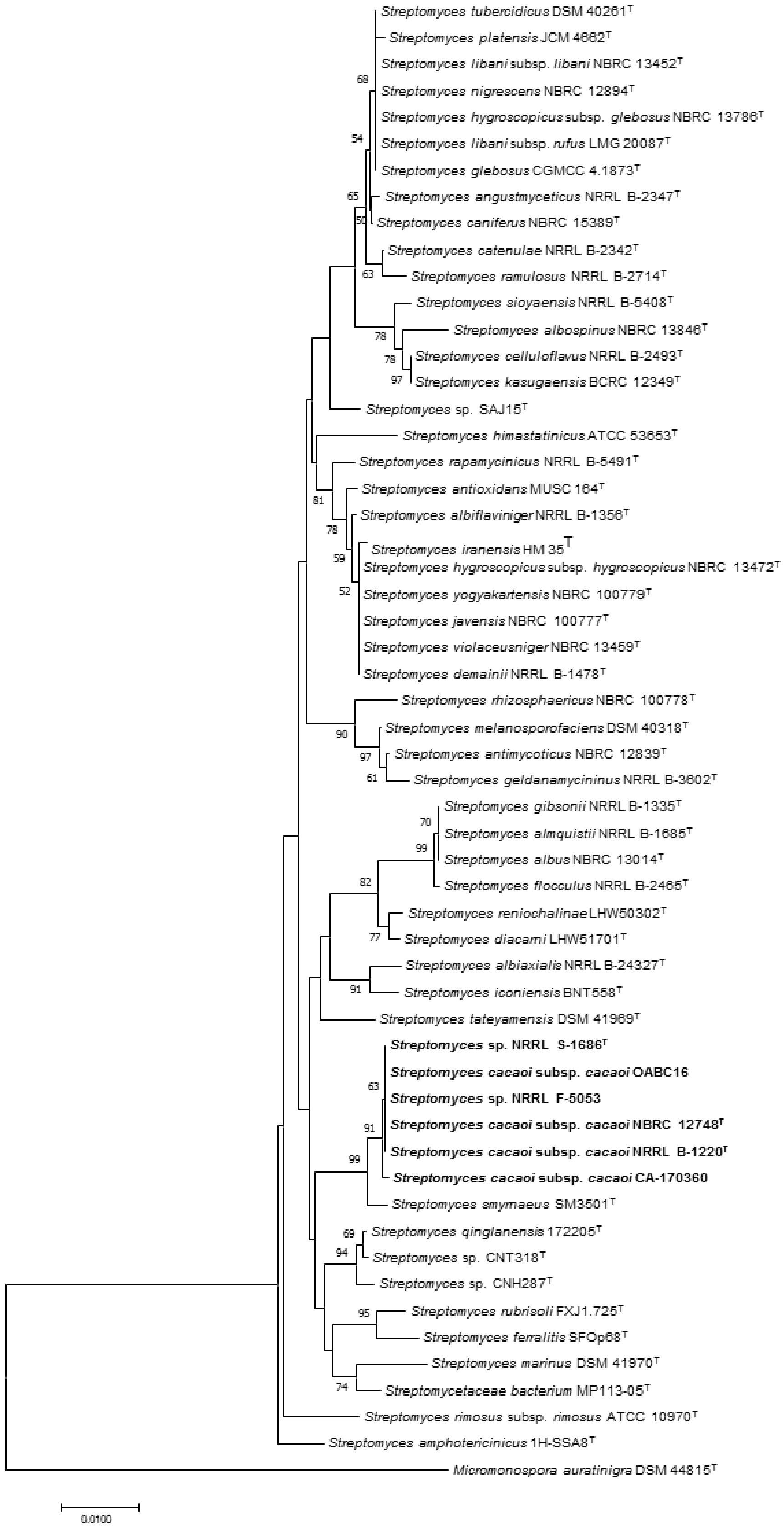
Streptomyces tubercidicus DSM 40261$^{T}$
Streptomyces platensis JCM 4662$^{T}$
Streptomyces libani subsp. libani NBRC 13452$^{T}$
68
Streptomyces nigrescens NBRC 12894$^{T}$
Streptomyces hygroscopicus subsp. glebosus NBRC 13786$^{T}$
54
Streptomyces libani subsp. rufus LMG 20087$^{T}$
Streptomyces glebosus CGMCC 4.1873$^{T}$
65
Streptomyces angustmyceticus NRRL B-2347$^{T}$
50
Streptomyces caniferus NBRC 15389$^{T}$
Streptomyces catenulae NRRL B-2342$^{T}$
63
Streptomyces ramulosus NRRL B-2714$^{T}$
Streptomyces sioyaensis NRRL B-5408$^{T}$
Streptomyces albospinus NBRC 13846$^{T}$
78
78
Streptomyces celluloflavus NRRL B-2493$^{T}$
97
Streptomyces kasugaensis BCRC 12349$^{T}$
Streptomyces sp. SAJ15$^{T}$
Streptomyces himastatinicus ATCC 53653$^{T}$
Streptomyces rapamycinicus NRRL B-5491$^{T}$
81
Streptomyces antioxidans MUSC 164$^{T}$
Streptomyces albiflaviniger NRRL B-1356$^{T}$
78
Streptomyces iranensis HM 35$^{T}$
59
Streptomyces hygroscopicus subsp. hygroscopicus NBRC 13472$^{T}$
52
Streptomyces yogyakartensis NBRC 100779$^{T}$
Streptomyces javensis NBRC 100777$^{T}$
Streptomyces violaceusniger NBRC 13459$^{T}$
Streptomyces demainii NRRL B-1478$^{T}$
Streptomyces rhizosphaericus NBRC 100778$^{T}$
Streptomyces melanosporofaciens DSM 40318$^{T}$
90
97
Streptomyces antimycoticus NBRC 12839$^{T}$
61
Streptomyces geldanamycininus NRRL B-3602$^{T}$
Streptomyces gibsonii NRRL B-1335$^{T}$
70
Streptomyces almquistii NRRL B-1685$^{T}$
99
Streptomyces albus NBRC 13014$^{T}$
82
Streptomyces flocculus NRRL B-2465$^{T}$
Streptomyces reniochalinae LHW50302$^{T}$
77
Streptomyces diacarni LHW51701$^{T}$
Streptomyces albiaxialis NRRL B-24327$^{T}$
91
Streptomyces iconiensis BNT558$^{T}$
Streptomyces tateyamensis DSM 41969$^{T}$
**Streptomyces sp. NRRL S-1686$^{T}$**
**Streptomyces cacaoi subsp. cacaoi OABC16**
63
**Streptomyces sp. NRRL F-5053**
91
**Streptomyces cacaoi subsp. cacaoi NBRC 12748$^{T}$**
99
**Streptomyces cacaoi subsp. cacaoi NRRL B-1220$^{T}$**
**Streptomyces cacaoi subsp. cacaoi CA-170360**
Streptomyces smyrnaeus SM3501$^{T}$
69
Streptomyces qinglanensis 172205$^{T}$
94
Streptomyces sp. CNT318$^{T}$
Streptomyces sp. CNH287$^{T}$
95
Streptomyces rubrisoli FXJ1.725$^{T}$
Streptomyces ferralitis SFOp68$^{T}$
Streptomyces marinus DSM 41970$^{T}$
74
Streptomycetaceae bacterium MP113-05$^{T}$
Streptomyces rimosus subsp. rimosus ATCC 10970$^{T}$
Streptomyces amphotericinicus 1H-SSA8$^{T}$
Micromonospora auratinigra DSM 44815$^{T}$
0.0100

*Supporting Figure 4. Neighbor-joining tree built with Mega X based on nearly complete 16S rRNA gene sequences of CA-170360 and the 50 closest type strains of the genus* Streptomyces*.* Micromonospora auratinigra *DSM 44815 (T) was used as an outgroup. The numbers at the nodes indicate the bootstrap value (%) based on NJ analysis of 1000 replicates; only values higher than 50% are shown.*

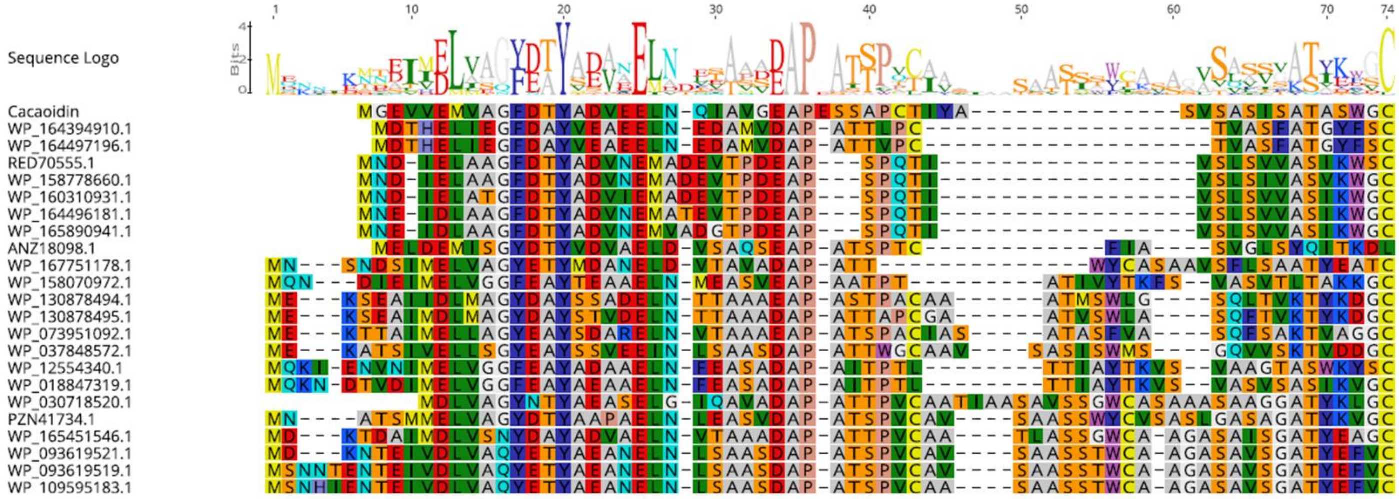


*Supporting Figure 5. Alignment of cacaoidin and putative lanthidins precursor peptides (accession numbers are shown). Sequence logo shows conserved residues.*

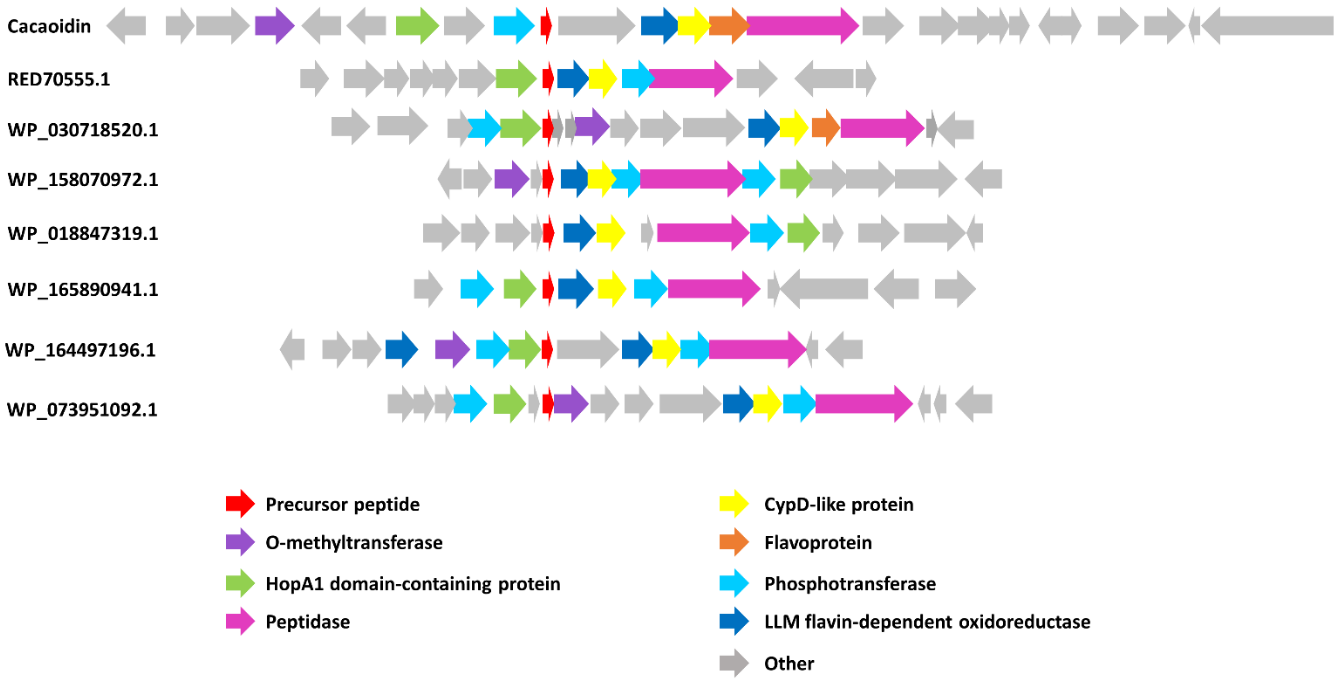


*Supporting Figure 6. Schematic representation of cacaoidin BGC and the BGC of some putative lanthidins. Accession numbers of each putative structural peptide are indicated.*

### References


1. Chater, K. F. and Wilde, L. C. (**1980**). Streptomyces albus G mutants defective in the SalGI restriction-modification system. *J. Gen. Microbiol*. 116, 323–334, DOI:10.1099/00221287-116-2-323

2. Zhang, J. J., Yamanaka, K., Tang, X. and Moore, B. S. (**2019**). Direct cloning and heterologous expression of natural product biosynthetic gene clusters by transformation-associated recombination. Methods Enzymol. 621, 87-110, DOI: 10.1016/bs.mie.2019.02.026

3. Kieser, T., Bibb, M. J., Buttner, M. J., Chater, K. F., and Hopwood, D. A. (**2000**). “Preparation and analysis of genomic and plasmid DNA,” in Practical Streptomyces Genetics, Vol. 1, eds T. Kieser, M. J. Bibb, M. J. Buttner, K. F. Chater, and D. A. Hopwood (Norwich: John Innes Foundation), 161–210.

4. Blin, K., Wolf, T., Chevrette, M. G., Lu, X., Schwalen, C. J., Kautsar, S. A., Suarez-Duran, H. G., de Los Santos, E. L. C., Kim, H. U., Nave, M., Dickschat, J. S., Mitchell, D. A., Shelest, E., Breitling, R., Takano, E., Lee, S. Y., Weber, T. and Medema, M. H. (**2017)** antiSMASH 4.0-improvements in chemistry prediction and gene cluster boundary identification. *Nucleic Acids Res*. 45(W1),W36-W41, DOI: 10.1093/nar/gkx319

5. van Heel, A. J., de Jong, A., Song, C., Viel, J. H., Kok, J. and Kuipers, O. P. (**2018**). BAGEL4: a user-friendly web server to thoroughly mine RiPPs and bacteriocins. *Nucleic Acids Res.* 46(W1), W278-W281, DOI: 10.1093/nar/gky383

6. Skinnider, M. A., Merwin, N. J., Johnston, C.W. and Magarvey, N. A. (**2017**). PRISM 3: Expanded prediction of natural product chemical structures from microbial genomes. *Nucleic Acids Res*. 45, W49–W54, DOI: 10.1093/nar/gkx320

7. Agrawal, P., Khater, S., Gupta, M., Sain, N. and Mohanty, D. (**2017**). RiPPMiner: a bioinformatics resource for deciphering chemical structures of RiPPs based on prediction of cleavage and cross-links. *Nucleic Acids Res*. 45, W80-W88, DOI: 10.1093/nar/gkx408

8. Johnson, M., Zaretskaya, I., Raytselis, Y., Merezhuk, Y., McGinnis, S. and Madden, T. L. (**2008**). NCBI BLAST: a better web interface. *Nucleic Acids Research*. 36, W5–W9, DOI: 10.1093/nar/gkn201

9. Zimmermann, L., Stephens, A., Nam, S. Z., Rau, D., Kübler, J., Lozajic, M., Gabler, F., Söding, J., Lupas, A. N. and Alva, V. (**2018**). A Completely Reimplemented MPI Bioinformatics Toolkit with a New HHpred Server at its Core. *J Mol Biol.* 2836(17), 30587-30589.

10. Jiang, W., Zhao, X., Gabrieli, T., Lou, C., Ebenstein, Y. and Zhu, T. F. (**2015**). Cas9-assisted targeting of chromosome segments CATCH enables one-step targeted cloning of large gene clusters. *Nat. Commun*. 6, 8101. DOI: 10.1038/ncomms9101

11. Jian, W. and Zhu, T. F. (**2016**). Targeted isolation and cloning of 100-kb microbial genomic sequences by Cas9-assisted targeting of chromosome segments. *Nat Protoc*. 11(5), 960-975, DOI: 10.1038/nprot.2016.055

12. Tong, Y., Robertsen, H. L., Blin, K., Weber, T. and Lee, S. Y. (**2018**). CRISPR-Cas9 Toolkit for Actinomycete Genome Editing. *Methods Mol. Biol*. 1671, 163-184, DOI: 10.1007/978-1-4939-7295-1_11
